# Intracellular TDP-43 amyloid nucleates from arrested nascent condensates

**DOI:** 10.64898/2026.02.10.705006

**Authors:** Jianzheng Wu, Shriram Venkatesan, Jacob Jensen, Tayla Miller, Jeffrey J. Lange, Sean McKinney, Einar Halldorsson, Zulin Yu, Vignesh Babu, Laura Sancho Salazar, Jeff Haug, Jay Unruh, Randal Halfmann

## Abstract

TDP-43 is a model protein for pathophysiological phase transitions, forming a multitude of intracellular assemblies with different physical properties. Physiological condensation is widely presumed to precede pathological aggregation, but the causal relationships between different modes of assembly in vivo are still unclear. Here we use Distributed Amphifluoric FRET (DAmFRET) and complementary approaches to map the phase space of TDP-43 self-assembly in yeast cells. We discovered that the low-complexity C-terminal domain (CTD) on its own partitions into soluble clusters that dynamically arrest en route to liquid-liquid phase separation. These clusters uniquely supported amyloid formation, and only when templated by pre-existing amyloids of other proteins. Self-interacting modules outside the CTD, whether in the full-length TDP-43, pathological C-terminal fragments, or fusion partners, all suppressed amyloid nucleation. They did this by promoting CTD condensation beyond the arrested state. Stress and cotranslational self-association had the same effect. We leveraged this property of condensation to stop CTD amyloid formation by co-expressing an oligomeric binder in cells. Our findings reveal that TDP-43 amyloid formation occurs only under very specific physical and biological circumstances that present new opportunities for therapeutic control.

## Introduction

Amyloid aggregates are effectively irreversible, and susceptible proteins are constantly driven toward that fate (Ciryam et al., 2015; Vecchi et al., 2020). Why then do amyloids take decades to form in patients?

TDP-43 is among the most frequent and problematic amyloids in the elderly, depositing in the neurons of approximately 1 in 5 people over age 80, where it contributes to ALS, Alzheimer’s Disease, and related dementias (Meneses et al., 2021; Nelson et al., 2019). It has consequently become a model for liquid-to-solid pathological phase transitions. The protein normally exists as diffuse multimers (Afroz et al., 2017; Fang et al., 2014; Sun et al., 2019; Zacco et al., 2022) that give way to dynamic condensates under stress (Maharana et al., 2018; Molliex et al., 2015; Schmidt and Rohatgi, 2016), and further to irreversible aggregates or amyloids with prolonged stress, aging, and disease progression (Arseni et al., 2023, 2022; Berning and Walker, 2019; Candelise et al., 2023; Cao et al., 2019; Chhangani et al., 2021; Gasset-Rosa et al., 2019; Lee et al., 2021; Lin et al., 2024; Nonaka et al., 2013; Shenoy et al., 2020; Tziortzouda et al., 2021). Our understanding of the physical bases of these transitions relies primarily on observations of purified protein under nonphysiological conditions and reaction volumes, and on microscopy of intracellular puncta containing thousands of subunits. However, amyloid formation is rate-limited by primary nucleation in the confined volumes of living cells, and pathological TDP-43 dysfunction precedes its visible aggregation (Chang et al., 2023; Spence et al., 2024). We consequently know very little about the underlying dynamics of nanoscopic multimers and the nature of the rate-limiting steps of TDP-43 amyloid formation in cells.

TDP-43 features an N-terminal homo-oligomerization domain (NTD), nuclear localization signal (NLS), two RNA recognition motifs (RRMs), and a C-terminal domain (CTD) of low sequence complexity (**Fig. 1A**). The largely disordered CTD is enriched in polar uncharged residues and a conserved hydrophobic patch, which forms transient secondary structure that is important to its self-association (Conicella et al., 2020, 2016; Gu et al., 2023; Jiang et al., 2013; Lim et al., 2016). A scattering of aromatic and aliphatic residues distributed throughout the CTD also contribute to phase separation (Mohanty et al., 2023; Schmidt et al., 2019). The many mutations that have been identified to accelerate disease onset and/or progression occur almost exclusively in the CTD. These are presumed to accelerate amyloid formation *in vivo*, because patient-derived aggregates are enriched for C-terminal fragments (CTFs) of TDP-43 (Feneberg et al., 2021; Forgrave et al., 2024), and the hydrophobic patch of the CTD dominates the ordered cores of those aggregates (Arseni et al., 2023, 2022).

**Figure 1.**
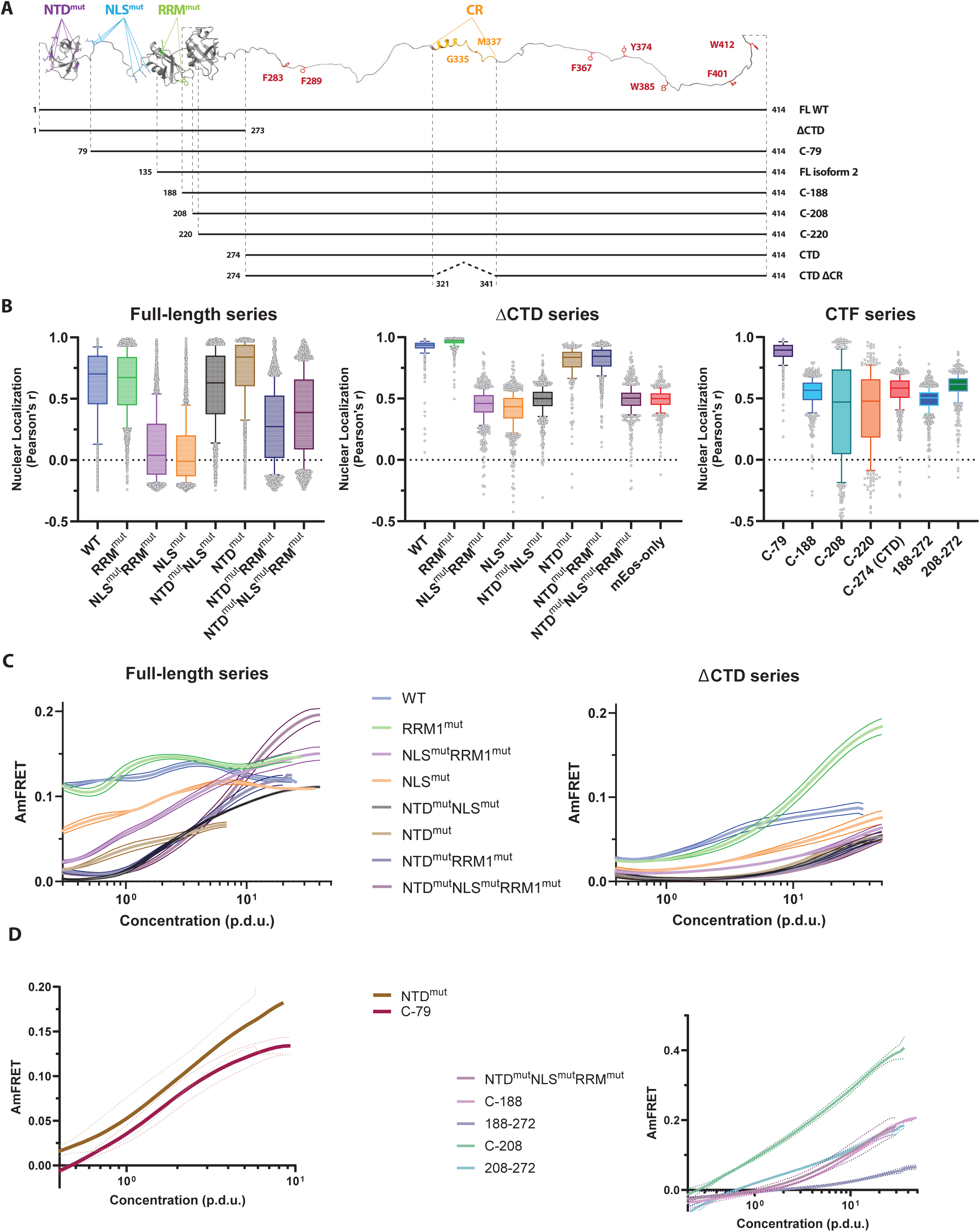
TDP-43 phase behavior is spatially regulated by the cooperative action of functional modules. A) **Top:** AlphaFold3-derived schematic of TDP-43, with relevant residues colored and labeled. NTD^mut^: E14A, E17A, E21A, R52A, R55A; NLS^mut^: K82A, K84A, K95A, K97A, R83A, R98A; RRM^mut^: W113A, R151A; key CR mutations: G335I, M337P, and Δ321-341. **Bottom:** Schema of fragments used to probe steric and functional contributions to phase behavior. All constructs have native C-termini and mEos-fused N-termini. B) **Comprehensive quantification of nuclear enrichment.** Pearson correlation coefficient between a nuclear marker and mEos fluorescence for the indicated fusions. The analysis highlights the synergy between the NTD and RRM in maintaining the nuclear pool of TDP-43, as well as the driving role of the CTD in cytoplasmic localization. See Methods for analysis details. Box represents median and IQR, and whiskers - 10^th^ & 90^th^ percentile from 1000s of single cell measurements. C) **Systematic DAmFRET analysis of domain variants** reveals the NTD’s role in native oligomerization, the triple mutant’s unmasking of concentration-dependent self-assembly, and the CTD’s role as the driver of condensation. Overlays of spline fits of median AmFRET versus derived concentration of the query protein. All fits include triplicate colony read-outs and trace the mean ± S.D. P.D.U., procedure-derived units normalized by C_trans_ of CTD WT (run-matched); detailed in Methods. The color key is identical for both series, with CTD-only (red curve) serving as the bridge sample between datasets. For example, RRM^mut^ in the left plot is in the context of the full-length protein, while in the right plot it represents the point mutations in the context of TDP-43 lacking the C-terminal domain entirely. Unless otherwise mentioned, AmFRET value of zero represents the median AmFRET of monomer-control post-compensation. D) **DAmFRET plots with spline fit overlays comparing CTFs to relevant point mutants.** The overt similarity between truncations and specific point mutants reveals that native N-terminal interactions, rather than steric bulk, modulate self-assembly. Conversely, the CTF C-208 exhibits aberrantly strong aggregation (much steeper DAmFRET slope), indicating a gain-of-function resulting from truncation of the otherwise-folded RRM2 domain. Plots represent multiple independent replicate experiments.

In addition to pathological amyloids, the CTD also drives physiological TDP-43 condensation, and this is widely seen as a prerequisite for amyloid formation (Babinchak and Surewicz, 2020; Mukherjee et al., 2024; Visser et al., 2024). The relationship of condensation to amyloid is likely to be nuanced, however, as recent in vitro and in silico studies have uncovered opposing roles of the interior and interface for the condensate-to-amyloid transition of proteins resembling TDP-43 (Alshareedah et al., 2024; Das et al., 2025; Emmanouilidis et al., 2024; Garaizar et al., 2022; Linsenmeier et al., 2023; Shen et al., 2023). Each of the functional modules of full-length (FL) protein have been found to impact the protein’s phase behavior in different ways (Chen et al., 2019; Maharana et al., 2018; Oiwa et al., 2023; Wang et al., 2018; Winton et al., 2008), but their combinatorial effects have not been studied systematically. Similarly, the different CTFs have very different aggregation propensities and contribute differentially to disease progression in animal models (Igaz et al., 2009; Kitamura et al., 2024; Yang et al., 2010). The most studied CTFs, known as “TDP-25”, have a gain of toxic activity associated with the misfolding and aggregation of truncated RRM2 (Berning and Walker, 2019; Kitamura et al., 2024; Kuo et al., 2009; Mackness et al., 2014; Morgan et al., 2017; Tavella et al., 2018; Wei et al., 2017). Nevertheless, CTFs beginning downstream of RRM2, i.e. just the CTD, appear to have the highest discriminatory power for TDP-43 pathology (Forgrave et al., 2024), suggesting that non-CTD-mediated aggregation as in TDP-25 may not translate to pathological amyloid formation.

Here we employed DAmFRET and complementary techniques across a panel of rational TDP-43 sequence variants to systematically map the contributions of TDP-43’s various functional modules to localization and phase behavior in living cells. Our results recapitulate known sequence determinants of the protein’s self-association and subcellular localization, while uncovering a previously unknown tendency of the CTD to dynamically arrest as nanoscopic clusters. Other oligomerizing modules, whether in the full-length protein, C-terminal fragments, or fluorescent protein fusions, allowed for further condensation, as did accelerated translation. In all cases, condensation potently inhibited amyloid nucleation. This property along with a massive conformational nucleation barrier dominate the kinetics of amyloid formation.

## Results

### Synergistic contributions to localization and phase behavior by TDP-43’s functional modules

To characterize TDP-43’s intracellular phase behavior systematically, we sought to express numerous variants of the protein under the same conditions across a range of concentrations spanning potential phase boundaries. Simultaneously, we needed to distinguish finite oligomerization, condensation, and amyloid-like aggregation. To achieve these goals, we rationally designed a library of sequence variants and cloned them as genetic fusions to mEos3.1 (hereafter, “mEos”). This exceptionally soluble, photoconvertible, and SDS-resistant fluorescent tag is uniquely suited to a powerful combination of assays: confocal fluorescence microscopy, distributed amphifluoric FRET (DAmFRET), and semidenaturing detergent-agarose gel electrophoresis (SDD-AGE) (Khan et al., 2018; Zhang et al., 2012). Microscopy allows direct visualization of large protein assemblies and subcellular localization; DAmFRET reports on size-independent protein assembly as a function of concentration; and SDD-AGE distinguishes assembly types by their size and stability.

Because the CTD of TDP-43 is the principal determinant of its phase behavior and multiple regulatory modifications occur very near the C-terminus, we fused mEos to the N-terminus of each variant to preserve the sterics and mobility of the native C-terminus. As the expression host we used a strain of budding yeast that we previously engineered to quantize amyloid aggregation events by halting cell division during exogenous protein expression. Except where indicated in subsequent sections, all experiments were conducted in the absence of endogenous amyloids (i.e. in [*pin^-^*] cells) that can accelerate amyloid nucleation by exogenous proteins. Consequently, and as will be shown in a subsequent section, puncta observed in this section are non-amyloid in nature.

We first assessed the concentration-dependence of subcellular localization and self-association of the WT full-length (FL) protein using high throughput confocal fluorescence microscopy and DAmFRET. We found that it localized predominantly to the nucleus (**Fig. 1B, S1A, B**), but with some cells exhibiting granular cytoplasmic fluorescence (**Fig. S1B**). Despite these different localization patterns, the protein assembled to approximately the same extent in all cells, as indicated by a uniform distribution of positive AmFRET values (“AmFRET” is the ratio of sensitized emission FRET intensity to direct-excited acceptor fluorophore intensity, and relates directly to the fractional extent and density of assemblies; **Fig. 1C, S1C**). The observed localization differences therefore appear to reflect underlying cell heterogeneities that are sensitive to TDP-43, rather than transitions in its material state. We also found that the protein restricted cell sizes, indicating a growth defect, resulting in a tightened distribution of concentrations relative to mEos expressed on its own (**Fig. S1D**).

We next assessed the contributions of each functional module of the FL protein to these behaviors. Given that even non-interacting linkers and spacers influence phase separation (Pappu et al., 2023), we first inactivated each functional module using rational point mutations rather than deletions to probe their contributions to phase behavior individually and combinatorially (**Fig. 1A**). Specifically, we disrupted homo-oligomerization of the NTD using interfacial mutants E14A, E17A, E21A, R52A, R55A (NTD^mut^; (Afroz et al., 2017)); nuclear localization using K82A, K84A, K95A, K97A, R83A, R98A (NLS^mut^; (Winton et al., 2008)); and nucleic acid binding using W113A, R151A (RRM^mut^; note that RRM2 does not directly bind nucleic acids (Ayala et al., 2005; Buratti and Baralle, 2001)).

The contributions of each module proved consistent with their expected functions. NTD^mut^ universally monomerized the protein at low concentrations while enhancing nuclear localization (**Fig. 1B, C, S1A, B, C**), confirming its predominant role in oligomerization by the FL protein. NLS^mut^ eliminated nuclear localization and concomitantly enhanced cytoplasmic foci (**Fig. 1B, S1A, B**). It also slightly reduced AmFRET (**Fig. 1C, S1C**), potentially as a simple consequence of the increased volume of the cytoplasm relative to the nucleus. RRM^mut^ enhanced nuclear localization in a manner that depended on the NLS (**Fig. 1B, S1A, B**). Combining RRM^mut^ with NTD^mut^ decreased AmFRET at low concentrations and alleviated the growth defect of the WT protein (**Fig. 1C, S1C, D**), suggesting its functional inactivation. Combining all three disruptions (NTD^mut^ NLS^mut^ RRM^mut^) caused a transition to high AmFRET at high concentrations (**Fig. 1C, S1C**), reminiscent of liquid-liquid phase separation previously characterized by DAmFRET (Khan et al., 2018; Miller et al., 2023). Altogether these data support an interpretation that integrates all modules into the major function of TDP-43 as an RNA-shuttling protein. Specifically, the protein oligomerizes in the nucleus through a combination of RNA binding and NTD-mediated interactions. The resulting mRNPs then exit the nucleus. Impeding either RNA-binding or NTD-oligomerization consequently strands TDP-43 in the nucleus. Pathological CTFs lack some or all of these modules depending on the N-terminal cleavage site. It is unclear to what extent the phase behavior of individual CTFs can be attributed to the loss of specific activities, the loss of steric bulk and consequent CTD exposure, or the insolubility associated with RRM2 cleavage and misfolding as in TDP-25. To evaluate all of these effects, we next designed and tested a series of synthetic CTFs (**Fig. 1A**). We found that a minimal truncation eliminating just the NTD (C-79) behaved indistinguishably from NTD^mut^, while a more extreme truncation eliminating the NTD, NLS, and RRM1 (C-188) resembled the triple disruptant (**Fig. 1D**), collectively revealing that steric changes contribute minimally to localization and aggregation. In marked contrast, CTFs with a slightly more extreme truncation including part of RRM2 (C-208 and C-220; representing TDP-25), or a previously uncharacterized splice isoform (2) that disrupts RRM1, aggregated profusely (**Fig. 1D, S1B, C**).

We next examined the contributions of the CTD by deleting it from the FL mutants. Deleting the CTD reduced AmFRET at low expression (**Fig. 1C, S1C**) and eliminated mislocalization to cytoplasmic puncta in all cases (**Fig. S1B**). It also eliminated the concentration-dependent transition to high AmFRET by the triple disruptant mutant (**Fig. 1C, S1C**), establishing that the CTD provides low affinity interactions that are necessary for condensation of FL protein oligomers in the cytoplasm. Interestingly, deleting the CTD in conjunction with RRM^mut^ caused the protein to transition to high AmFRET at high concentrations, in a manner that depended on the WT NTD and NLS (**Fig. 1C, S1C**). Hence, RNA-binding and the CTD restrict the extent of NTD-driven assembly in the nucleus.

### TDP-43 CTD forms dynamically arrested soluble clusters

We next characterized the CTD itself. We found that it transitioned from low to high AmFRET in the low micromolar regime (estimated from prior standardization of DAmFRET data (Khan et al., 2018); **Fig. 2A**), in agreement with saturating concentrations of CTD observed in vitro (Conicella et al., 2016, 2020; Li et al., 2018b; Schmidt and Rohatgi, 2016). Given prior reports of condensation of the CTD in cells, and the similarity of its DAmFRET profile with that of the puncta-forming triple-disrupted FL protein, we expected the same. Surprisingly, however, the protein remained fully diffuse in the majority of cells irrespective of its expression level when imaged by confocal microscopy (**Fig. 2B**). Given the lack of a clear phase boundary, we will refer to the approximate concentration required for cooperative self-assembly as the transition concentration (C_trans_; **Fig. S2A**).

**Figure 2.**
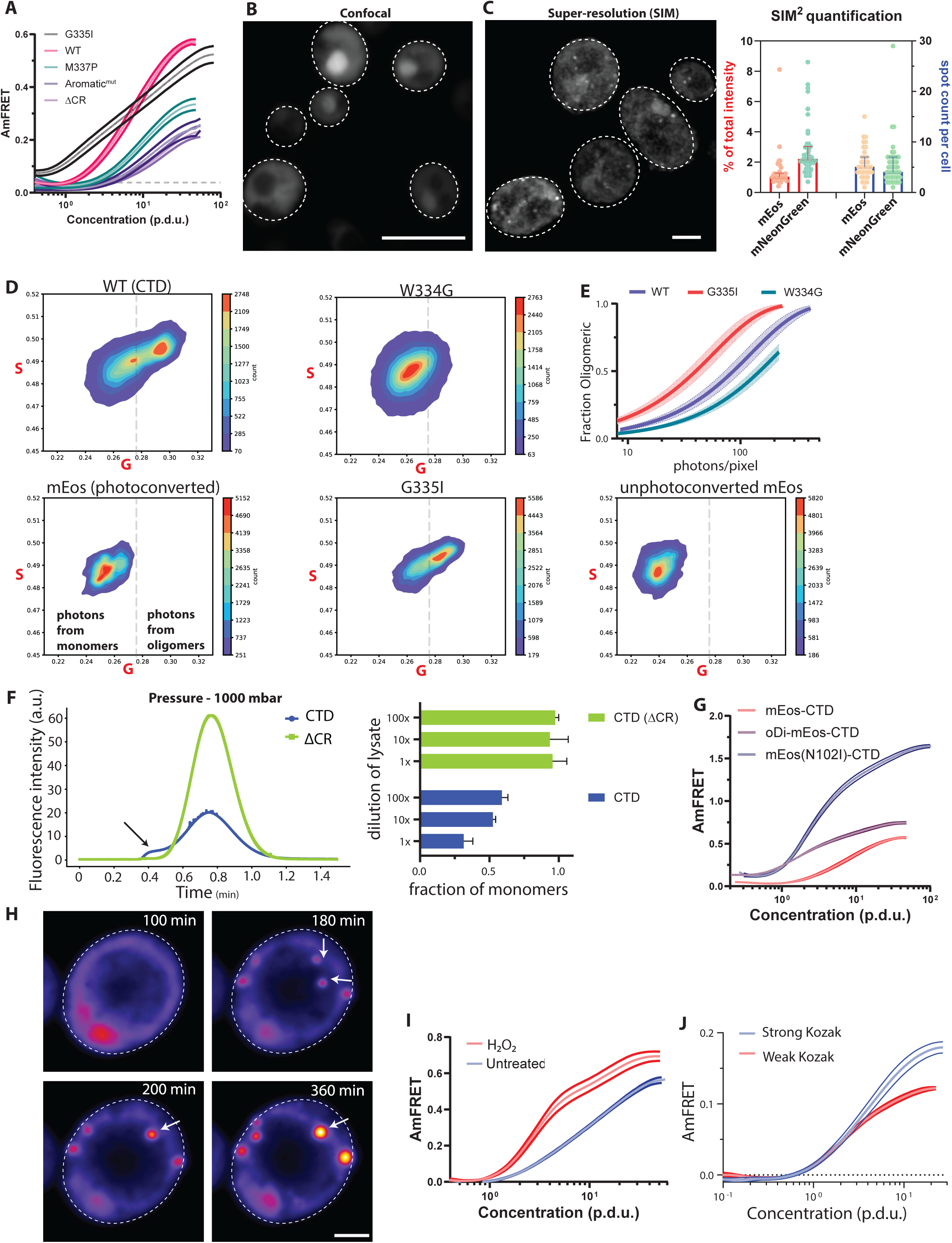
TDP-43 CTD forms dynamically arrested clusters regulated by translation flux and oxidative stress. A) TDP-43 CTD self-assembles cooperatively, modulated by sequence determinants of canonical phase separation: Spline overlay of WT CTD exhibiting a sharp transition to high AmFRET values at a specific concentration (C_trans_) and documented mutations shifting C_trans_ as expected. “Aromatic ^mut^” denotes mutations of the following residues to A (alanine) - F289, 367 & 401; Y374; W385 & 412. The horizontal dashed line shows AmFRET levels of mEos by itself. Plotted are Mean ± S.D. of triplicates. B) **CTD resists condensation** even when expressed 50-fold beyond C_trans_. A max-projected confocally imaged field of cells expressing mEos-tagged CTD showing entirely diffuse fluorescence. Bright accumulations are the nucleus. Scale bar = 10 μm. C) **CTD assembles into sub-diffraction clusters, not droplets:** Left: Representative field from 70 nm resolution structured illumination microscopy (SIM) of mNeonGreen-tagged CTD revealing the presence of a few diffraction limited spots at expression levels approximately 50-fold higher than C_trans_. Scale bar = 2 μm. Right: mEos-tagged CTD shows similar results. Bar height = Median; Error bars = IQR. D) FLIM-FRET phasor plots of WT and solubilizing mutant (W334G) of mEos-CTD. G and S, the cosine and sine components of the raw lifetime trace at the instrument’s repetition rate. Dashed line drawn at a constant x-intercept to visually demarcate the two populations on the plot. E) Analysis of FLIM-FRET data showing the fraction of photons in the oligomeric population as a function of pixel intensity, for WT, W334G, and G335I. Lines show fits to one phase association ± 95% CI. F) **Biophysical sizing of soluble oligomers.** (Left): Flow Induced Dispersion Analysis (FIDA) of clarified lysates of CTD at 1000 mbar mobilization pressure reveals a distinct subpopulation of multimers (black arrowhead) of R_h_ 5-50nm only when the CR is intact. Plot representative of at least triplicates. (Right): Derived monomeric fraction of serially diluted clarified lysates of CTD- and CTD (ΔCR)-expressing yeast, revealing CR-dependent persistent multimeric species (5-50 nm range) at 1000 mbar mobilization pressure, even after dilution up to 100-fold for 6 hours. Shown are means ± S.D. of triplicates. G) **Cluster growth is valence-limited:** A dimerizing mutation to mEos (N102I) and fusion of CTD to a homodimer, oDi, induce macromolecular condensation with minimal change in transition concentration. Spline fits ± S.D. of triplicate DAmFRET profiles. H) **Oxidative stress triggers CTD coalescence:** Representative montage of time lapse imaging of CTD-expressing yeast upon treatment with 3 mM H_2_O_2_ monitored for 6 hours, showing the emergence of round puncta (**top, right**), that fuse (**bottom, left**) and continue growing (**bottom, right**) indicating liquid behavior. Images processed in Fiji and false colored with Fire scheme LUT. I) Spline fitting of DAmFRET profiles of the H_2_O_2_ treated population of CTD-expressing cells reveals a significant increase in cooperativity beyond C_trans_, without affecting C_trans_, suggesting it increases surface tension without strengthening the major interactions. CTD-expressing cells were treated with an 8 mM dose for 2 hours before DAmFRET. J) **Altering translation flux of CTD specifically reveals kinetic control of clustering.** A pair of Kozak sequences designed to initiate translation strongly and weakly, showing that cluster growth occurs cotranslationally, as seen by decreased AmFRET steady state levels in the spline fit with weak Kozak initiation context. Shown are means ± S.D. of triplicates.

Because we did not observe visible condensates of mEos-tagged CTD by confocal microscopy, we resorted to super-resolution imaging of PFA-fixed cells (to circumvent diffusion-mediated blurring) using Structured Illumination Microscopy (SIM). In addition to mEos-CTD, we tagged the CTD with mNeonGreen, one of the brightest and most photostable monomeric fluorescent proteins. At an estimated 70-80 nm resolution, we found that the brightest cells had on average five diffraction limited spots. However, the spots only accounted for 1-2% of the total fluorescence per cell (**Fig. 2C**). The steep increase in AmFRET beyond C_trans_, combined with expression levels reaching 50-fold excess, demonstrates that the diffuse pool is predominantly multimeric species smaller than 70 nm.

Other studies of TDP-43 CTD have reported condensation in unstressed cells, albeit using different fluorescent tags (e.g. (Johnson et al., 2008; Nonaka et al., 2009)). To explore consequences of fluorescent protein choice to CTD phase separation, we surveyed puncta formation using alternative fluorescent proteins. CTD remained diffuse when fused to mEos, mNeonGreen, mCherry, or mScarlet-I, but formed puncta when fused to mEGFP, SYFP2, or mKelly2 (**Fig. S2B**). Hence, CTD is prone to condense aberrantly when fused to certain fluorescent proteins, a phenomenon that it shares with other IDRs (Barkley et al., 2024; Dörner et al., 2025; Fatti et al., 2025).

To explore the physical basis of CTD clusters, we mutated sequence features that had previously been shown to govern CTD phase separation *in vitro*, and measured the corresponding changes with DAmFRET. Specifically, we either increased (G335I) or decreased (M337P) intermolecular CR interactions, deleted the entire CR segment (Δ321-341), or ablated aromatic side chains outside the CR (F283A, F289A, F367A, Y374A, W385A, F401A, and W412A or “Aromatic^mut^”). These mutations each produced the expected decrease (G335I) or increase (others) in C_trans_ (**Fig. 2A, S2C**), confirming that subresolution clusters and phase separation are driven by similar multivalent interactions. The dependence on multivalency and the occasional incidence of visible puncta of different sizes suggests that clustering results from dynamical arrest of nascent condensates (Pappu et al., 2023; Ranganathan and Shakhnovich, 2020). Dynamical arrest occurs from an internal saturation of binding sites within a cluster, leaving the surface unable to recruit more molecules (Pappu et al., 2023; Ranganathan and Shakhnovich, 2020; Tan et al., 2025).

To quantify the fraction of CTD molecules partitioning into clusters, we modified a confocal microscope to measure the fluorescence lifetimes of excited-state mEos molecules. Fluorescence lifetimes are sensitive to the proximity of the molecules via FRET -- which we enhanced through partial photoconversion (FLIM-FRET) -- but are insensitive to photobleaching and concentration. Hence, different lifetime populations correspond to different protein states, and the relative sizes of the different populations reflect the composition of protein states in each imaged pixel. We analyzed yeast cells expressing either mEos alone, WT CTD, or CTD bearing mutations that either enhance (G335I) or disrupt (W334G) CR interactions. We confirmed that mEos alone and the W334G CTD mutant produced a single largely overlapping population of photon lifetimes, representing the monomeric protein (**Fig. 2D**). In contrast, WT and G335I variants produced both the monomeric population of photon lifetimes and a second population with longer lifetimes consistent with multimers. The fraction of pixels in this population increased with fluorescence intensity, and did so at lower intensities for G335I than for WT (**Fig. 2E**), confirming that the former associates more strongly than the latter.

As an orthogonal measure of soluble clusters, we used flow-induced dispersion analysis (FIDA) to evaluate the hydrodynamic radii (R_h_) and stability of fluorescent particles in clarified crude lysates. We first captured data at conditions that can resolve R_h_ of multimers less than 50 nm, and found that at expression levels below C_trans_, mEos-CTD diffused as expected for a monomer (Table 1, methods). In contrast, at the normal range of expression levels for DAmFRET (which span C_trans_), the protein diffused as a mixture of monomers and oligomers evidenced by higher R_h_. We next raised the capillary pressure to push particles larger than 5 nm outside of Taylor conditions (Chamieh et al., 2017; Taylor, 1953). This produced a shoulder of fluorescence of slower diffusing molecules that dispersed asymmetrically (though consequently, R_h_ could no longer be determined directly), validating the existence of polydisperse multimers in the size range of 5-50 nm (**Fig. 2F**). Deleting the CR eliminated these species and largely restored monomeric diffusion (**Fig. 2F**). Finally, we diluted the CTD-expressing lysate up to 100-fold with lysis buffer, incubated for 6 hrs to allow for dissociation of multimers, and repeated the analysis. The multimers persisted (**Fig. 2F**), revealing vastly slower dissolution than has been described for CTD droplets that form with similar threshold concentrations (Conicella et al., 2016). The multimers therefore harden or “mature” after formation. Importantly, this behavior of CTD is not an artifact of our fluorescent protein fusion, as dye-labeled CTD similarly formed stable “nanocondensates” under near-physiological conditions in vitro (Houx et al., 2024).

**Table 1.** Radii of hydration (Rh) in yeast lysate.

| protein | observed Rh ± SD (nm) | n | expected Rh ± SD (nm) |
| --- | --- | --- | --- |
| mEos-CTD, with expression below C<sub>trans</sub> | 3.809 ± 0.078 | 7 | 3.956 ± 0.009 |
| mEos-CTD | 4.528 ± 0.069 | 5 | 3.956 ± 0.009 |
| mEos-CTD G335I | 4.693 ± 0.042 | 6 | 3.973 ± 0.010 |
| mEos-CTD ΔCR | 3.247 ± 0.031 | 3 | 3.834 ± 0.009 |
| mEos | 2.903 ± 0.0115 | 3 | 2.596 ± 0.001 |

To test if cluster growth is indeed valence-limited, we introduced a point mutation to mEos (N102I) to restore the dimerizing tendency of its progenitor (Zhang et al., 2012). This change increased AmFRET at all concentrations (**Fig. 2G, S2D**) and allowed CTD to form large round puncta (**Fig. S2E**). The mutation nevertheless did not lower C_trans_ beyond the two-fold (1.91 ± 0.24) reduction expected from dimerization of the molecule, confirming that clustering itself is driven by cooperative interactions in the CTD, while effectively doubling those interactions suffices to allow coalescence. In addition, we fused mEos-CTD to a homodimeric coiled coil (oDi) as described previously (Fletcher et al., 2012; Kandola et al., 2023), and found that it qualitatively phenocopied mEos (N102I) (**Fig 2G, S2D, E**).

Cytoplasmic TDP-43 condensation can be induced by oxidative stressors such as hydrogen peroxide (H_2_O_2_) (Cohen et al., 2015; Gasset-Rosa et al., 2019; Zuo et al., 2021). To explore the influence of stress on dynamical arrest, we exposed yeast cells expressing mEos-CTD overnight to 3 mM H_2_O_2_ and took time lapse images for up to 6 hours. H_2_O_2_ induced CTD LLPS in this context, as indicated by the appearance of liquid-like puncta that continued to grow following fusion (**Fig. 2H**, **S2F**). This coincided with a sharp rise in AmFRET beyond C_trans_, even though the value of C_trans_ itself changed little (p > 0.05, T-test, paired, two-tailed) (**Fig. 2I, S2G**). H_2_O_2_ therefore allows clusters to coalesce without overtly influencing their principal interactions, as if by specifically softening their shells. This could happen, for example, through direct oxidation of certain CTD methionines (Gu et al., 2023; Lin et al., 2020; Mohanty et al., 2023; Ozguney et al., 2025) and/or reduced chaperone binding (Carrasco et al., 2023).

In cells, the high density of nascent protein at polysomes promotes homo-oligomerization and condensation (Bertolini et al., 2021; Heidenreich et al., 2020; He et al., 2025). This effect may counteract dynamical arrest, thus promoting cluster growth. To explore this possibility, we manipulated the rate of translation initiation by mutating the Kozak sequence (Park and Subramaniam, 2019) of the CTD ORF and qualitatively validated it in our system (**Fig. S2H**). We found that specifically decelerating translation flux of the construct reduced AmFRET levels as well as H_2_O_2_-mediated condensation irrespective of CTD concentration (**Fig. 2J, S2I**), thereby definitively establishing that CTD condensation is under dynamical control in cells.

### CTD amyloid nucleates specifically from arrested clusters

TDP-43 remains soluble for decades in human neurons. Amyloid formation in the tiny volumes of living cells is rate-limited by a nucleating conformational change that can be accelerated by the presence of other amyloids (Derkatch et al., 2001; Keefer et al., 2017; Khan et al., 2018). Therefore, to study the emergence of amyloid in the context of arrested clusters, we next expressed CTD in cells containing an innocuous self-perpetuating amyloid form ([*PIN*^+^]) of the low-abundance endogenous protein Rnq1, which are genetically identical to the [*pin^-^*] cells used above. To verify that amyloid had formed in the [*PIN*^+^] cells, we extracted the detergent-resistant insoluble fraction of whole cell lysates, revealing a single band corresponding to the expected molecular weight of mEos-CTD (**Fig. S3A**). No bands were observed in cells that did not express mEos-CTD.

We then used DAmFRET to distinguish cells that contained CTD amyloid from those that did not, based on the expected gain in AmFRET for amyloid due to its higher density and depletion of monomeric protein. The DAmFRET profile of CTD-expressing [*PIN*^+^] cells closely resembled that of their [*pin*^-^] counterparts, including the presence of monomer-only and cluster-containing cells demarcated by C_trans_, but with a key exception. The [*PIN^+^*] cells contained a third and fourth population each with distinctly higher AmFRET (**Fig. 3A**). These populations were discontinuous yet overlapping with the cluster-containing population. The discontinuities indicate true phase transitions, and the overlap indicates that the transitions are not determined solely by concentration on the timescale of our experiment. These transitions are therefore subject to large kinetic barriers rooted in the extreme conformational changes required for amyloid nucleation (Khan et al., 2018).

**Figure 3.**
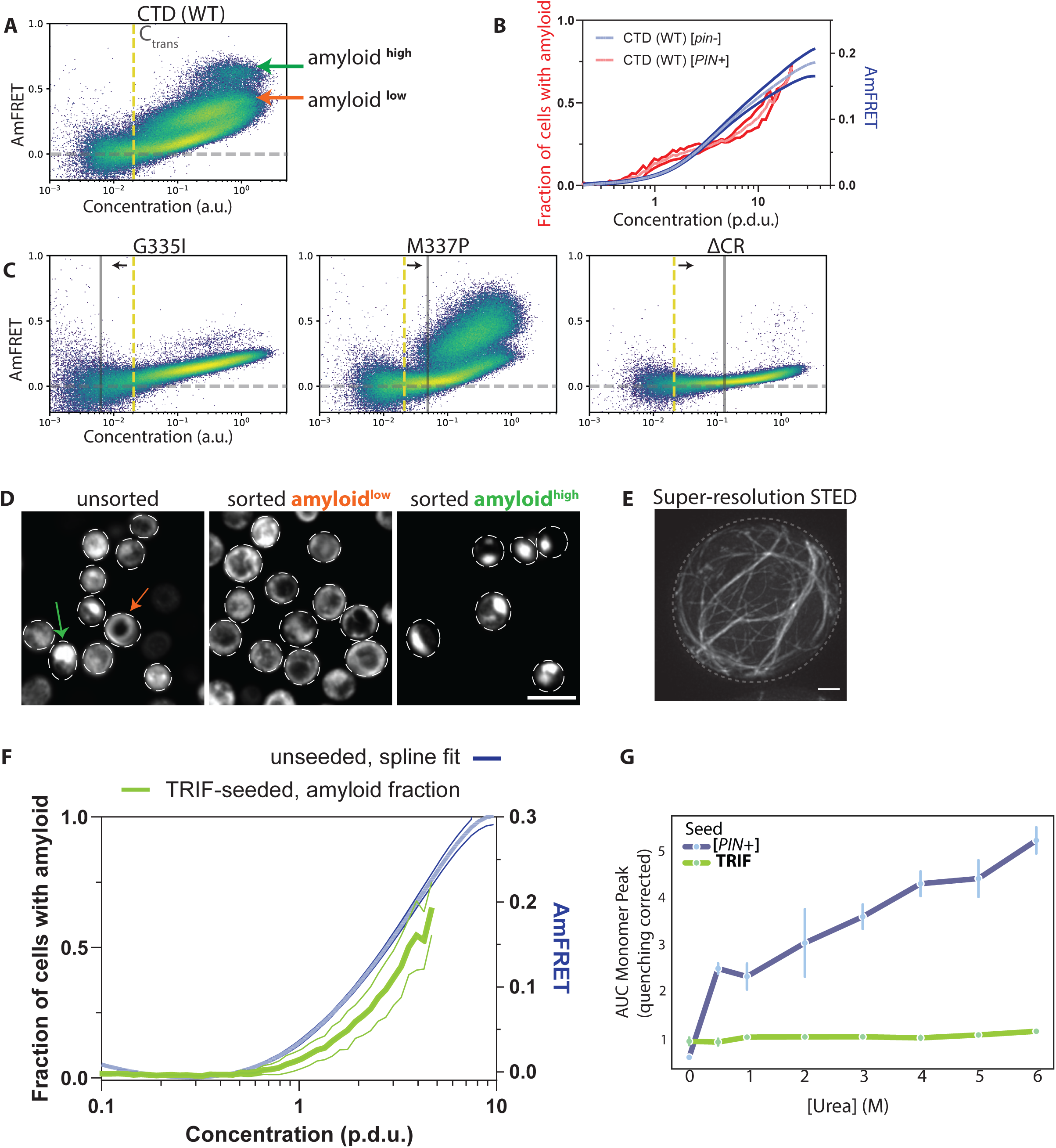
Dynamically arrested clustering is required for CTD amyloid formation. A) **Amyloid nucleation coincides with clusters.** CTD forms amyloid exclusively in yeast containing the endogenous [*PIN*^+^] amyloid. DAmFRET reveals key attributes of TDP-43 amyloid - density plot shows that CTD exhibits two discontinuous higher AmFRET populations (denoted by orange and green arrows) while the cluster-containing population (the sloped region) persists from [*pin*^-^] cells, indicating a kinetic barrier to the formation of amyloid from arrested clusters.The dashed yellow line indicates the C_trans_ of wild-type CTD (WT) derived from run-matched [*pin-*] cells (non-amyloid condition). B) **Complex concentration dependence of nucleation.** Binned DAmFRET plot and clustering of the populations to account for the fraction of cells in the discontinuous higher-AmFRET populations versus the rest reveals the concentration dependence of CTD amyloid. Overlaying the spline fit of AmFRET vs concentration (p.d.u.) from run-matched [*pin^-^*] profiles reveals that nucleation begins sharply at the transition concentration (C_trans_), suggesting that clusters are a prerequisite for CTD amyloid. The subsequent plateau suggests that cluster growth or factor saturation limits nucleation before a second permissive regime emerges at higher concentrations. Shown are means ± S.D. of triplicates. C) **Mutations strengthen the connection between the two species of CTD.** DAmFRET analysis of CTD mutants in [*PIN^+^*] cells shows that the concentration onset of amyloid shifts in concordance with the specific C_trans_ of each mutant (e.g., lower for G335I, higher for M337P), reinforcing the dependence of nucleation on the initial clustered state. Deletion of the CR nearly eliminates amyloid, in accordance with its involvement in the amyloid core. Solid grey line maps the C_trans_ derived from [*pin^-^*] run-matched cells. Black arrows indicate the direction of change of C_trans_ relative to WT CTD (denoted in shaded yellow line). D) **Morphology tracks with amyloid “type”.** Single confocal fluorescent slices of CTD amyloid-containing cells sorted by their DAmFRET signature using spectral FACS. Cells in the amyloid^low^ population contain fibrillar-looking aggregates even in dim cells, whereas amyloid^high^ cells contain prominent bright amorphous puncta with a near absence of diffuse fluorescence, occurring exclusively at high protein concentration. Differently colored arrows point to the two morphologies in the micrographs. Scale bar = 10 μm. E) A representative cell from STED super-resolution imaging of mNeonGreen-tagged CTD amyloid (in [*PIN^+^*]), upon deconvolution (elaborated in Methods), reveals a network of fibrils. Scale bar = 1 μm. F) **Cross-seeding by a human amyloid-forming protein.** DAmFRET analysis of CTD in yeast expressing human TRIF (a functional amyloid implicated in ALS) instead of Rnq1 showing cross-seeding of CTD amyloid. **(Above)**: Overlay of unseeded AmFRET spline and tracing of amyloid-enrichment over concentration shows CTD amyloid’s dependence on multimeric clustering even when seeded by TRIF. **(Below)**: Histogram overlay of the seeded populations in triplicates shows TRIF preferentially induces an amyloid^high^ state. G) **Template identity determines polymorph stability.** Urea denaturation curves of CTD amyloids derived from resolving monomer and amyloid fractions by FIDA reveals the resistance of amyloid, from lysates of the respective populations of cells, to Urea denaturation. Amyloids nucleated by TRIF are significantly more stable than those nucleated by [*PIN^+^*], confirming that polymorph selection is governed kinetically.

Plotting the fraction of cells with amyloid as a function of CTD concentration revealed a complex concentration-dependence with multiple inflections (**Fig. 3B)**. Amyloid failed to form at concentrations lower than C_trans_. The fraction of amyloid-containing cells rose sharply around C_trans_ -- suggesting clusters promote nucleation -- before plateauing briefly and then increasing again at very high concentrations. We interpret the plateau as evidence that clusters lose potential for amyloid formation as they grow, either by acquiring nonpermissive conformations (Lin et al., 2024; Shuster and Lee, 2022) or saturating an amyloid-enabling cellular factor. Finally, the resumption of amyloid formation at very high concentrations reveals a second pathway for amyloid formation involving either higher-order clustering (albeit still without obvious phase separation) or saturation of an amyloid-*inhibiting* cellular factor.

To examine the potential role of clusters in amyloid formation, we compared the concentration-dependence of amyloid formation for the aforementioned CTD mutants that alter C_trans_. The concentration at which amyloid began to form tracked with C_trans_ (**Fig. 3C**): lower for G335I and higher for M337P. This suggests that the clusters are on-pathway to amyloid. Unexpectedly, however, the kinetic barrier to amyloid formation beyond C_trans_ showed the opposite relationship: a much smaller fraction of cells formed amyloid for G335I than for M337P (**Fig. 3C, S3B, D**). Deleting the helical region greatly reduced amyloid formation, and again only beyond (the greatly increased) C_trans_, as expected from its participation in the fibrillar cores of the available pathologic TDP-43 amyloid structures (Arseni et al., 2024, 2023, 2022). Intriguingly, the two amyloidogenic regimes of CTD concentration corresponded to the two distinct populations of amyloid-containing cells, one with higher AmFRET than the other (**Fig. 3A**). The low-AmFRET type (amyloid^low^) formed at and just beyond C_trans_, whereas the high-AmFRET type (amyloid^high^) formed at much higher concentrations. Fluorescence microscopy upon spectral FACS sorting revealed that the amyloid^low^ cells contained fibrillar aggregates, while amyloid^high^ cells instead contained giant amorphous puncta (**Fig. 3D, S3C**). To observe the filamentous morphology of the more abundant amyloid^low^ state, we used STED microscopy to evaluate mNeonGreen-CTD subcellular localization and aggregate morphology in [*PIN*^+^] cells, revealing cytoplasmic ribbon-like bundles of filaments characteristic of amyloids (**Fig. 3E**).

Given the strict dependence of CTD amyloid formation on a pre-existing conformational template, we wondered if the identity of the template mattered. As Rnq1 is not conserved in humans, we specifically asked if CTD amyloid formation can also cross-seed off of endogenous human amyloids. To do so, we deleted *RNQ1* and expressed human TRIF, an intracellular amyloid in pro-inflammatory signaling that has been functionally linked to ALS pathogenesis (Baker et al., 2022; Gentle et al., 2017; Komine et al., 2018; Richardson et al., 2023). CTD amyloid formation was restored in the presence of TRIF (**Fig. 3F**), and it again only occurred beyond C_trans_, suggesting that clusters are critical for amyloid nucleation even for a physiological cross-seeding template. Interestingly, CTD preferentially formed an amyloid^high^ state when cross-seeded by TRIF (**Fig. S3E**).

To determine if the two amyloids states have underlying structural differences, we incubated lysates of [*PIN*^+^]- and TRIF-templated CTD with varying concentrations of urea and used FIDA to measure the amount of protein solubilized by each treatment. The resulting dissolution curves (**Fig 3G, S3F**) reveal that TRIF-templated amyloids (predominantly amyloid^high^) are much more stable than [*PIN*^+^]-templated amyloids (predominantly amyloid^low^), confirming that the distinct AmFRET populations correspond to different CTD amyloid structures, or polymorphs, and that the preferred polymorph can be determined by pre-existing amyloids in the cell.

### Condensation inhibits amyloid formation

Having revealed that CTD amyloid can form from soluble clusters, but that the clusters become recalcitrant to amyloid formation as total concentration increases, we next explored the impact of phase separation on amyloid formation. To do so, we first subjected our full panel of TDP-43 FL and CTF variants to DAmFRET in [*PIN*^+^] cells. The resulting profiles (**Fig. S4A**) largely resembled their counterparts in [*pin*^-^] cells, showing that [*PIN^+^*] amyloids do not overtly influence TDP-43’s non-amyloid interactions. In fact, WT, RRM^mut^, and C-208 behaved the same in both contexts, suggesting a total absence of amyloid formation (**Fig. 4A, S4A, C**). However, all FL variants with NTD^mut^ and/or NLS^mut^ yielded a dusting of higher AmFRET cells (**Fig. 4A, S4A**), and C-188 populated a discrete high AmFRET population closely resembling that of CTD itself (**Fig. S4C**). In all cases, deleting the CTD eliminated the high-AmFRET population, consistent with its having resulted from amyloid (**Fig. S4B**). The fact that the NTD and NLS each suppress amyloid formation is intriguing because they have opposite effects on nuclear localization. However, both mutations increased solubility at low concentrations, unlike RRM^mut^ which neither increased solubility nor allowed for amyloid. Similarly, the least soluble variant, C-208, did not form amyloid (**Fig. S4C**). Collectively, these data strongly suggest that TDP-43 amyloid nucleates in the solution phase.

**Figure 4.**
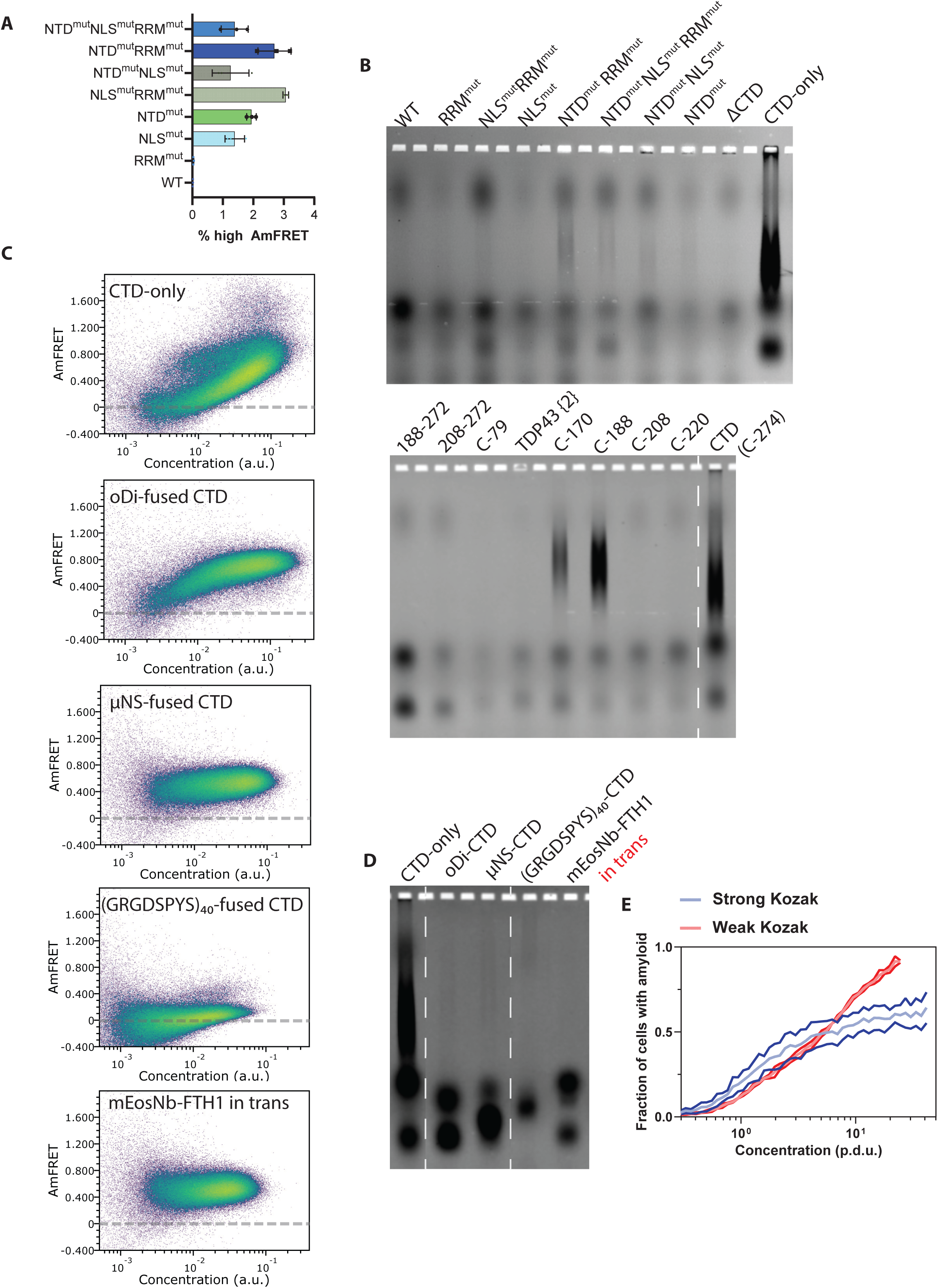
Physiological condensation suppresses amyloid nucleation. A) **Native domains suppress amyloid formation.** High-AmFRET populations were quantified by gating against run-matched [*pin^-^*] controls to exclude non-amyloid assemblies. While TDP-43 proved recalcitrant to amyloid formation, overall, full-length variants with disrupted oligomerization (NTD^mut^) or localization (NLS^mut^) populate a high-AmFRET tail, indicating that native interactions protect against amyloid conversion. B) **Biochemical confirmation of amyloid.** SDD-AGE analysis of detergent-insoluble complexes confirms DAmFRET interpretation that WT TDP-43 and C-208 lack the characteristic high-molecular-weight smear of amyloid, and validated the discontinuous DAmFRET profile of C-188 as amyloid. CTD-alone formed a smear while WT protein devoid of CTD did not, underscoring the necessity of CTD to form amyloids of TDP-43. Dashed white line demarcates splices within the same gel, while solid white line demarcates splicing between gels. C) **Induced condensation of TDP-43 CTD blocks its amyloid formation.** Panel of a diverse set of condensate-inducing fusions to CTD - μNS, (GRGDSPYS)_40_, and mEos (N102I) - all diminish CTD amyloid in [*PIN*^+^] in the timescale of the DAmFRET experiment. D) SDD-AGE of the constructs in C), confirming reduced amyloid smears by condensate-forming fusions relative to CTD alone. E) **Reducing cluster growth promotes amyloid.** Decelerating translation flux using a weak Kozak sequence prevents the plateau in amyloid formation at high concentrations (red trace). Shown are means ± S.D. of triplicates.

The small AmFRET differences between cells containing presumed amyloid and non-amyloid self-assemblies necessitated an orthogonal assay for amyloid formation. For this purpose we used semidenaturing detergent-agarose gel electrophoresis (SDD-AGE), which resolves detergent (SDS)-insoluble complexes by size. Amyloids are polydisperse high molecular weight complexes that resist dissolution by SDS, yielding a characteristic smear with SDD-AGE. We found that the FL WT protein, C-208, and splice isoform 2 lacked a smear in both [*pin*^-^] and [*PIN^+^*] cells, while C-188 and the CTD itself formed a smear specifically in [*PIN^+^*] cells (**Fig. 4B, S4D**). The SDD-AGE data therefore confirm the interpretations from DAmFRET, collectively demonstrating that the intrinsic amyloid propensity of the CTD is masked by other modules of the FL protein and by the RRM2 fragment in TDP-25.

Our findings thus far counter the prevailing paradigm that liquid-liquid phase separation precedes and accelerates amyloid formation by TDP-43. Does phase separation instead inhibit amyloid formation in cells? To answer this question, we used several genetic approaches to induce CTD condensation and measured the resulting effects on amyloid formation. First, we appended either of two well-characterized condensate-forming sequences to the N-terminus of mEos in the CTD fusion proteins: the self-assembling coiled coil region from the virus protein µNS (Gama et al., 2024; Holliday et al., 2019; Rodriguez Gama et al., 2022; Schmitz et al., 2009), and a resilin-inspired intrinsically disordered sequence, (GRGDSPYS)_40_ (Dzuricky et al., 2020). Second, we induced CTD condensation by dimerizing mEos as described above with oDi or by coexpressing an FTH1-fused nanobody against mEos in trans with mEos-CTD (Bracha et al., 2018; Kandola et al., 2023; Kimbrough et al., 2025b). We expressed the constructs in [*PIN*^+^] cells, wherein they all formed large puncta as expected (**Fig. S4E**). We then characterized these cells with DAmFRET and discovered that all four changes greatly reduced or eliminated discontinuous high-FRET populations of cells (**Fig. 4C**). Similarly, using SDD-AGE, we observed a severe reduction in SDS-resistant high-molecular weight species (**Fig. 4D**). These results confirm that the amyloidogenic nature of CTD oligomers is tempered by condensation, whether artificially or through the functional activities of other domains in the native context of full-length TDP-43.

Finally, to probe the relationship of amyloid nucleation to the extent of condensate progression prior to dynamical arrest, and independently of any change to the protein itself, we reduced translation flux using the aforementioned Kozak mutation. Remarkably, this change greatly increased the fraction of amyloid containing cells beyond C_trans_ for WT CTD, and partially rescued the amyloid formation defect of the G335I mutant (**Fig. 4E, S4F**). Hence, even though amyloid nucleation strictly requires CTD multimerization, it is impeded by growth or coarsening of those multimers en route to condensates.

## Discussion

We systematically surveyed the contributions to phase behavior and subcellular localization by each of the functional modules of TDP-43, leading to an integrated model of functional TDP-43 oligomerization and condensation. Not surprisingly, both condensation and amyloid formation depended on the CTD. However, all changes that influenced condensation of the CTD, whether to other modules of the FL protein, to the CTD itself, or to the fluorescent protein to which it was fused, oppositely impacted amyloid formation beyond C_trans_.

Amyloid nucleation by CTD within the timescale of our experiments strictly depended on a cross-seeding template. This template apparently can have multiple identities, including endogenous TRIF amyloids that amplify programmed cell death signaling in human cells. These or other physiological or pathological amyloids plausibly accelerate TDP-43 proteopathy in vivo. The specific amyloid structure (polymorph) of TDP-43 defines distinct diseases (Scheres et al., 2023). While the differences between amyloid structures of TDP-43 may be determined by their different pathological contexts, our findings show that they can, alternatively, be determined by the cross-seeding template in otherwise identical cells. This phenomenon, which may be common among polymorphic amyloids (Sharma and Liebman, 2013; Yuzu et al., 2021), suggests a potential mechanistic basis for the inception of different TDP-43 proteopathies, whereby the amyloid structure established upon primary nucleation predetermines the course of the disease.

Our results lead to a model whereby amyloid nucleation occurs specifically at the surfaces of dynamically arrested clusters, a nanoscopic intermediate that precedes true phase separation. This model is strongly supported by several lines of evidence: (1) CTD multimers have a heavy-tailed size distribution with occasional visible puncta; (2) once formed, they persist after dilution, indicating a kinetically stable, “hardened” state distinct from classic LLPS; (3) their assembly involves multivalent interactions yet becomes valence-limited, as adding a “sticker” outside the CTD circumvented arrest; (4) their extent of assembly is governed by translation flux, suggesting that growth competes with hardening. The behavior of CTD in cells bears some resemblance to that of purified Fus, which was recently reported to form stable arrested nanoclusters below the C_sat_ for condensation (Ge et al., 2025).

Our intracellular findings complement in vitro observations of soluble CTD multimers (Babinchak et al., 2019; Conicella et al., 2016; Houx et al., 2024; Li et al., 2018a), as well as recent in vitro and in silico observations pointing to a common tendency of amyloid to nucleate at the (solidified) surfaces of condensates (Bauer and Nikoubashman, 2024; Emmanouilidis et al., 2024; Espejo et al., 2025; Farag et al., 2022; Garaizar et al., 2022; Linsenmeier et al., 2023; Mahendran et al., 2025b; Shen et al., 2023; Tan et al., 2025; Zhang et al., 2025). The particular susceptibility of CTD to this phenomenon may stem from its detergent-like patchy distribution of hydrophobic and hydrophilic residues that tend to segregate to the condensate interior and interface, respectively (Zhang et al., 2025). The resulting micelle-like structure may facilitate the alignment of polypeptide backbones leading to an accumulation of beta structure at the interface (Bauer and Nikoubashman, 2024; Emmanouilidis et al., 2024; Espejo et al., 2025; Farag et al., 2022; Garaizar et al., 2022; Mahendran et al., 2025b; Shen et al., 2023; Tan et al., 2025; Zhang et al., 2025). Whether amyloid nucleation results from a specific conformational bias at the interface, or simply the increased ratio of surface area to volume in small clusters, remains to be determined.

Our integrated model also explains the natural suppression of CTD amyloid in the context of the FL protein. Non-amyloid oligomerization driven by the NTD, nuclear localization by the NLS, and RNA-binding by RRM1, all increased the extent of assembly at low concentrations and collectively eliminated amyloid formation. The NTD provides stickers that are completely orthogonal to the CTD, driving the protein past the small-cluster stage and into larger, non-amyloidogenic particles. In this way, these modules of the FL protein function as integral kinetic suppressors of amyloid, effectively accelerating coalescence and reducing the lifespan of the amyloid-prone intermediate.

Our data reveal that clustering and therefore the decision between physiological condensation and pathogenic amyloid is kinetically controlled by the cell. Specifically, reducing translation flux by mutating the Kozak sequence greatly increased the fraction of amyloid-containing cells. In essence, the faster the translation initiation rate, the greater the density of nascent CTD chains, and the more likely they are to escape dynamical arrest and subsequent amyloid formation. This otherwise paradoxical finding implies that cluster growth toward condensation is subject to a universal stress-responsive control parameter, and underscores the potential for therapeutic strategies to mitigate pathogenic aggregation through subtle tweaks to protein synthesis.

Many questions remain unanswered. Chief among them: how do the numerous pathogenic mutations in CTD influence amyloid formation? In the context of our model, they may not simply influence C_trans_. They could accelerate dynamical arrest, thereby increasing the population of amyloid-prone species; decrease the activation energy for the amyloid-nucleating conformation within the dynamically arrested clusters; or facilitate interactions with yet-to-be identified cellular templates. Exploring these possibilities will require systematic comparisons in a cellular context.

Similarly, multiple post-translational modifications (beyond proteolytic cleavage) such as ubiquitination and CTD phosphorylation are closely associated with TDP-43 pathology but remain to be studied in the context of dynamical arrest. CTD phosphorylation suppresses both condensation and amyloid formation (Gruijs da Silva et al., 2022) despite occurring outside the known structural determinants of either. In light of our findings, this observation suggests that phosphorylation may influence interfacial properties independently of the primary drivers of C_trans_ and directly impact amyloid competence.

Stress-induced phase separation is widely believed to trigger pathogenic aggregation by TDP-43 and related proteins (Gasset-Rosa et al., 2019; Li et al., 2013; Yan et al., 2025). Our findings join those of others (Das et al., 2025; Glineburg et al., 2024; Mahendran et al., 2025a; Wallace et al., 2015) in suggesting a counternarrative of critical therapeutic significance: that coalescence is a protective, anti-amyloid pathway. We here show in living cells that TDP-43 CTD tagged with minimal interference, forms finite arrested species rather than archetypal condensates. The reduction of interfacial surface area resulting from progression to droplets limits the nucleation of essentially irreversible amyloid aggregates. Therefore, rather than focusing on strategies to prevent TDP-43 clustering or condensation altogether (the prevailing approach), a more targeted and effective strategy may be to chemically or genetically accelerate the coalescence of clusters to effectively “bypass” the amyloidogenic intermediate.

## Acknowledgements

We thank Lucinda E. Maddera and Amanda Kroesen for assistance with imaging and data processing, Scott McCroskey for high-throughput yeast transformations, Megan Halfmann and Jeevapani Hettige for assistance with figure preparation. This work was funded by the National Institute Of General Medical Sciences of the National Institutes of Health (NIH) under Award Number R01GM130927 (to RH); the NIH through the Alzheimer’s Disease Center at the University of Kansas Medical Center (KUMC; grant P30 AG035982); the KU Endowment through a Woodyard Fellowship from the Institute for Neurological Discoveries at KUMC (to JGW), and the Stowers Institute for Medical Research. Original data underlying this manuscript can be accessed from the Stowers Original Data Repository at https://www.stowers.org/research/publications/LIBPB-2608.

## Methods

### Plasmid and yeast strain construction

Yeast strains rhy1713 and rhy1852 were used for most experiments (Khan et al., 2018). To visualize and quantify nuclear localization, we constructed rhy3239b in three steps as follows. The P2A ribosome skipping motif was deleted from vector CX (Miller et al., 2023) to produce vector DB, which served as a PCR template for linker-BDFP1.6:1.6 upon amplification with oligonucleotides (forward: 5’-ggtactagggctgttaccaaatactcctcctctactcaagccGGTGACGGTGCTGGTTTA-3’; reverse: 5’-aaagaaaacatgactaaatcacaatacctagtgagtgacttaTCGATGAATTCGAGCTCG-3’) containing homology (lowercase) to the 5’ end of the *HTB2* ORF. This PCR product was transformed into rhy2054 (Kimbrough et al., 2025c) to integrate via homologous recombination linker-BDFP1.6:1.6 prior to the stop codon in the endogenous *HTB2* ORF. This was followed by backcrossing to restore *ATG8*. Finally, the strain was converted to [*pin^-^*] by serial passaging on YPD plates containing 3 mM guanidine hydrochloride (GdHCl). Proteins of interest were expressed from the *GAL1* promoter in high-copy plasmids derived from V08 and V12 as described (Khan et al., 2018). See **Table S1** for plasmids including ORF sequences used in this study. Details are available upon request.

### DAmFRET

As previously described (Kandola et al., 2023; Kimbrough et al., 2025c).

### SDD-AGE

As previously described (Kandola et al., 2023).

### High content confocal microscopy

Replicate transformant colonies of yeast expressing TDP-43 query constructs were imaged using a high content microscope (Opera Phenix) as previously described (Kimbrough et al., 2025c) except using a PerkinElmer 40x water objective lens (N.A. 1.1). Images were captured for four fields and five confocal slices, typically amounting to hundreds of cells per well. For imaging the various fluorophore tags of TDP-43 CTD as in Fig. S2C, mEos3.1, mNeonGreen, EGFP and SYFP2 were captured using Ex/Em of 488/ (500-550) nm, and mKelly2, mScarlet-I and mCherry using Ex/Em of 561/ (570-630) nm laser-detector pairs.

### Quantification of nuclear localization

Nuclear enrichment of mEos-tagged TDP-43 and associated variants was assessed by quantifying the correlation of mEos intensity to BDFP1.6:1.6, in cells expressing HTB2 tagged to BDFP1.6:1.6, as a proxy for the nucleus. Cells were segmented on a SLURM cluster using Cellpose (Stringer and Pachitariu, 2025) and the boundaries eroded to avoid membrane edge effects. Pearson’s correlation coefficient from pixel data was calculated for each cell (an individual ROI). Wells with fewer than 50 cells were excluded from further analysis. The log-transformed mean fluorescence per cell followed a bimodal distribution with expressing cells clearly distinguishable from non-expressing. Non-expressing cells were excluded from further analysis. Pearson correlation was computed on individual masked cells using pearson_corr_coeff from scikit-image. Data from four fields was then consolidated and single cell values were plotted onto a box-whisker plot. Segmentation and quantification pipelines are available at https://github.com/jouyun/2026_WuVenkatesan.

### Super Resolution imaging of TDP-43 CTD non-amyloid multimers

Super-resolution imaging was performed on a ZEISS Elyra 7 Lattice SIM microscope equipped with a 63x oil immersion objective lens (Plan-Apochromat 63x/1.40 Oil). Z-stacks were acquired with a step size of 0.1 µm over 15 phases per slice. Raw images were reconstructed using ZEN software (black edition) with the “best fit” parameter. Processed images were generated using either standard SIM processing (sharpness setting of 9-10) or the SIM² deconvolution algorithm (under fixed setting). The SIM² method is based on the Richardson–Lucy iterative deconvolution and was applied to reduce artifacts and improve resolution.

Cells were segmented in 2D using Cellpose (cyto3 model). For spot detection, images underwent white top-hat filtering (50-pixel radius) and intensity normalization within each cell mask. Spots were then identified via an *h*-maxima transform (noise tolerance 0.4). For 3D quantification, raw Z-stacks were background-subtracted. To determine each cell’s Z-range, the mean Z-profile within the 2D cell mask was threshold. Total cell integrated intensity was calculated by summing all voxels within this defined 3D volume. To localize spots in 3D, the Z-profile within a 5-pixel lateral radius of each 2D spot was fitted to a 1D Gaussian to identify the precise sub-voxel Z-position. Spot integrated intensity was summed within this 5-pixel radius over a 7-slice Z-window (±3 slices) to cover the range of the entire spots. Finally, the total number of spots and the ratio of aggregate spot intensity to total cell intensity were quantified per cell.

### STED microscopy

STED images were acquired as described (Tu et al., 2023) on a Leica SP8 Gated STEDMicroscopy with 100×, 1.4 oil N.A. objective. Green channel (mNeonGreen) was excited with a pulsed white light (80 MHz) tuned to 488 nm and was depleted with a pulsed 592 nm laser at 80 to 90% maximum output. All STED images were acquired in 2D mode to maximize lateral resolution, and each image averaged eight times in the line average mode. Emission photons were collected with internal Leica HyD hybrid detectors with a time gate setting at 1 to 6 ns.

For images of amyloid-containing cells (Fig. 3E), raw STED data were processed post-acquisition by deconvolution using Huygens Professional (version 14.10) with a theoretical point spread function (PSF). Background intensity levels were manually sampled from the raw data, and signal-to-noise ratios (SNR) were maintained between 15 and 20 during processing.

### H_2_O_2_ timelapse

Timelapse images of protein induction in yeast cells were imaged on a Nikon Spinning Disc microscope built on a Ti base coupled to a spinning disc head (Yokogawa CSU W1). In this strain TDP43 was tagged with EGFP and excited with a 488nm laser through a 60x Plan Apochromat (NA 1.4) oil objective. The fluorescence was collected back through the same objective, through a bandpass filter (ET525/36m) onto an sCMOS camera (Hamamatsu Flash 4). Images were acquired in z stack format every 5 minutes for 4 hours with 11 z slices 0.5 microns apart. Transmitted light was also acquired. The laser power was set to 100% while the camera was adjusted to 50ms for a well saturated image. During the timelapse, cells were trapped and incubated in a CellASIC Onix device (Millipore). Cells were trapped in the diploid yeast chip, with fresh media flowing through the device at room temperature. In this case, we added 3 mM of H_2_O_2_ to the media at the start of imaging. After the data was collected, cells of interest were visualized using data processing steps in Fiji (https://fiji.sc/) with custom in house written plugins. Individual timelapse files were first concatenated together, then sample drift was corrected using “StackRegJ”. Individual cells were then cropped out of the full field of view. The Z stacks were then SUM projected through the entire z range. We verified manually that any puncta of interest were contained in the same z slice before sum projecting. Images were then scaled to 500×500 pixels with bilinear interpolation. In the montage, the brightness and contrast of each image was adjusted individually to improve visualization.

### Calculating concentration-dependence of amyloid

Concentration (a.u.) was determined as the ratio of compensated acceptor intensity and side scatter (SSC-A), a proxy for effective cell volume, as described previously (Kimbrough et al., 2025a; Miller et al., 2023). Concentration was binned into 64 logarithmically spaced bins, either within a range containing values for replicates, or the 0.1^th^ percentile and maximum value of Concentration for a single plot. Within each bin, the fraction of events in a high AmFRET population was calculated. Instead of mEos-only as the negative control, given the atypical profile of TDP-43 CTD, we chose run-matched non-amyloid, respective [*pin*^-^] plot for the CTD variant. The negative gates were defined as before (Kandola et al., 2023), but with upper gate values as either the 95^th^, 97^th^, or 98^th^ percentile based on the noise in the reference ([*pin^-^*]) and the [*PIN^+^*] plots.

### Spline fitting and determination of C_trans_

To visualize the relationship between AmFRET and protein concentration, data were partitioned into 64 concentration bins (a.u.). A “spline” was constructed by smoothing the median AmFRET values across these bins. To ensure statistical robustness, bins were pruned to include only those containing data from at least two of the three biological replicates. The initial low-concentration bins were excluded where signal fell below the assay’s limit of detection. Because absolute fluorescence intensities (a.u.) fluctuate between cytometer runs, the analyzed concentration range was chosen to match the experimentally relevant dynamic range. Final plots represent the mean of the triplicates, with error bars indicating the standard deviation. C_trans_ was determined similarly to C_sat_ in a prior study (Kimbrough et al., 2025a; Rodriguez Gama et al., 2026), except that here we used the maximum of the third derivative of AmFRET versus concentration.

### Longitudinal Data Acquisition and Normalization Strategy

Because datasets were generated over a span of several years by multiple experimentalists, raw fluorescence readouts exhibit batch effects. These arise from minor discrepancies in user-preferred cytometer settings and unavoidable hardware fluctuations, such as standard photoconversion lamp replacements (which, despite standardized energy output targets, can yield minor variations in total photon delivery).

To ensure robust cross-comparison, we employed a strict run-matched experimental design; no mutant or experimental condition was analyzed without a concurrently illuminated and acquired WT control. The AmFRET parameter, by virtue of its being ratiometric FRET (FRET-A/ Acceptor-A from raw cytometer read-outs for every single cell), tends to fluctuate across datasets. Consequently, AmFRET values are only directly comparable within a given figure panel but not across all panels, unless explicitly stated. Representative raw data plots display the native output settings of their specific acquisition run to maintain data integrity. For all derived analyses, plots, and multi-experiment quantifications, outputs were normalized against the C_trans_ of the run-matched WT CTD, shown as “Concentration (p.d.u.)” where p.d.u. stands for procedure derived units. This internal normalization unifies the scales across the study, allowing for accurate, internally controlled comparisons between panels regardless of the original acquisition parameters.

### Fluorescence Lifetime Imaging Microscopy - Fluorescence Resonance Energy Transfer (FLIM-FRET)

Images were acquired on a Leica SP8 microscope equipped with a pulsed white light laser, a DMi8 stand, and a FLIM module. Regular confocal images of pre-photoconverted mEos cells were excited with 488 and 561nm lasers lines for the green and red forms, respectively. Images were excited and collected through an HC Plan Apo 100x oil objective (NA 1.4). The fluorescence between 500-550nm was collected for the green form and between 611-700nm for the red form. Both images were acquired with different HyD detectors with 1024 x 1024 pixels. The FLIM data was then acquired for the same cells in the same field of view with the same pixel density with 16 line repetitions. Green mEos was excited with a pulsed 488nm laser line and collected through the same 500-550nm filter onto a HyD detector in FLIM mode. FLIM data was exported as individual bin files for further processing in Python.

### FLIM-FRET Image Analysis

Individual cells were segmented from summed fluorescence intensity images using Cellpose (‘cyto’ model) (Stringer et al., 2021). Cell masks were eroded to exclude boundary pixels susceptible to cross-cell contamination. Hand annotated nuclear masks were applied to classify each pixel as nuclear or cytoplasmic. Only pixels with total photon counts exceeding 8 were retained for downstream analysis. For phasor analysis (Ranjit et al., 2018), images were downsampled to 0.18um resolution and smoothed with a two-dimensional Gaussian filter (σ = 4.0 pixels) prior to analysis.

A phase correction was applied to account for the instrument response function. For each pixel, the phasor coordinates G and S were calculated as the first-harmonic cosine and sine transforms of the normalized fluorescence decay. To quantify the relative contribution of two fluorescence lifetime states, a linear decomposition was performed along a line connecting two reference phasor positions (G = 0.255, S = 0.485 and G = 0.295, S = 0.497). The fractional position along this line (FracB) was computed for each pixel using a dot-product projection, where FracB = 0 corresponds to the lower-FRET reference state and FracB = 1 corresponds to the higher-FRET state. Unmixing was performed by decomposing each pixel’s total fluorescence into two channels weighted by FracA (1 − FracB) and FracB, yielding two-channel images representing the spatial distribution of each lifetime state. All analyses were performed in Python using NumPy, SciPy, scikit-image, OpenCV, pandas, Cellpose, and napari.

### Yeast lysis for biochemical and FIDA measurements

To isolate amyloid aggregates from yeast, a combination of cryogenic grinding and detergent-based fractionation was employed. All steps were performed on ice or at 4°C unless otherwise specified to maintain protein stability.

The lysis buffer consisted of 50 mM HEPES (pH 7.5), 200 mM NaCl, 1% (v/v) Triton X-100, 1 mM EDTA, and 0.5% (v/v) Tween-20. Immediately prior to use, the buffer was supplemented with 4x Halt Protease and Phosphatase Inhibitor Cocktail, 0.5 mM DTT, and one cOmplete Protease Inhibitor Cocktail tablet per 50 mL.

Yeast cell pellets harvested from 200 mL cultures inoculated in SD-Ura overnight at 30C shaker incubator, followed by induction of expression of the respective TDP-43 construct driven by *GAL1* promoter, SRaff+Gal medium expressing the TDP-43 in unseeded yeast [*pin*^-^] or seeded [*PIN*^+^]/TRIF co-expressing cultures were processed as follows. Pellets were ground to a fine powder using a mortar and pestle pre-chilled with liquid nitrogen in equal volume of lysis buffer. Constant freezing was maintained by the periodic addition of liquid nitrogen throughout the grinding cycles. The resulting powder was transferred to pre-chilled tubes, centrifuges at 13000g for 10 min before using the supernatant fraction for measuring Rh values.

### Amyloid Enrichment

For amyloid enrichment, the powdery lysate was resuspended in 25 mL of the lysis buffer and suspension was incubated on ice for 10 minutes, with thorough mixing in between. Initial cell debris was removed by centrifugation at 2000g for 5 minutes. The supernatant was collected and further treated with N-lauroylsarcosine (sarcosyl) to a final concentration of 4% (w/v) or 1% SDS. Following a 10-minute incubation on ice, the sample was centrifuged at 12000g for 15 minutes to remove larger insoluble complexes. The detergent-soluble supernatants was transferred to Ti45-compatible ultracentrifuge tubes. To ensure rotor stability, volumes were adjusted to at least 75% capacity using Amyloid Isolation Buffer supplemented with 4% sarcosyl. Amyloid fibrils were pelleted via ultracentrifugation at 144,000g for 1 hour at 4°C. The resulting pellet was resuspended in 500 µL of the lysis buffer. To track extraction efficiency and protein distribution, aliquots from each fraction (total lysate, supernatants, and pellets) were collected and analyzed via SDS-PAGE.

### Thermodynamic stability of TDP-43 amyloids from lysate

Chemical depolymerization and thermodynamic stability of differentially seeded TDP-43 CTD amyloid was determined using urea-induced disassembly coupled to Flow-Induced Dispersion Analysis (FIDA) of the soluble fraction (FidaBio). Yeast cultures expressing the respective amyloid-seed combination were lysed and incubated for 2 hours at 25 °C in increasing concentrations of urea (0–6 M) prepared in HEPES-based lysis buffer + HALT protease inhibitor + 0.5 mM TCEP, with exactly 5 ul of lysate in each well and 15 ul of concentrated urea (4.3rd of its respective final concentration - 15 ul of 8M urea diluting to 6M final at 20 ul including lysate fraction). Following incubation, samples were analyzed by FIDA to quantify the residual monomer fraction. The analysis was performed using the method below (Table 2) with a 100cm length and 75µm inner diameter capillary and excited using a 480 nm LED. The method involved the following steps: washing using 1M NaOH (3500 mbar, 30 sec) and milliQ water (3500 mbar, 30 sec); equilibration using lysis buffer (3500 mbar, 20 sec); sample (75 mbar, 20 sec); and measurement using lysis buffer (1000 mbar, 90 sec) - all at 25 ℃. To account for fluorophore quenching by Urea, data normalized to that of yeast lysate expressing mEos-only for respective Urea concentration. Due to the overnight run time, samples were ordered in descending concentration of urea, with triplicates staggered, so as to normalize for any time dependent effects of amyloid stability across samples. For each elution profile, the fraction of soluble monomer (f) was calculated as the ratio of the area under the curve (AUC) corresponding to the monomer peak divided by the total integrated AUC for that sample. Data analysis was adopted from elsewhere (Farzadfard et al., 2024; Konstantoulea et al., 2025). This value reflects the proportion of protein that remains in the soluble state under each denaturant condition. The fit of all data points was performed by nonlinear least-squares minimization using the model function in lmfit (v1.3.4).

### Radius of hydration (Rh) estimation

We estimated the expected Rh using minimum dissipation approximation as in (Waszkiewicz et al., 2024). Using glm_mda_diffusion.ipynb, we set mEos as globular and allowed the rest of each sequence to be disordered. We set temperature at 298.15 K, viscosity at 0.8891 mPa⋅s, and used an ensemble size of 1000 with 100 bootstrap rounds.

### Cell sorting of TDP-43 amyloid populations

Cell sorting and fluorescence resonance energy transfer (FRET) analysis were performed on a BigFoot spectral cell sorter (Thermo Fisher Scientific) equipped with a 70 µm nozzle and operated with phosphate-buffered saline (PBS) as sheath fluid. mEosGreen (donor) was excited with a 488 nm laser, and donor emission was collected on the 507 nm PMT. A compensation of ∼8% was applied to correct donor spillover into the 583 nm PMT. mEosRed (acceptor) was excited with a 561 nm laser, and acceptor emission was collected on a separate 583 nm PMT. FRET was quantified as mEosRed emission in the 583 nm channel during 488 nm excitation (sensitized emission), whereas direct acceptor fluorescence was measured as 583 nm emission during 561 nm excitation. This configuration allows separate measurement of FRET and direct acceptor signals despite their overlapping emission spectra. Cells were first gated on forward and side scatter to identify cellular events, and singlets were selected by forward scatter area (FSC-A) versus height (FSC-H). Subsequent analysis was restricted to singlet events positive for mEosGreen fluorescence. Final sort gates were defined on a two-parameter plot of acceptor fluorescence (x-axis) versus FRET signal (y-axis).

---

**Figure S1.**
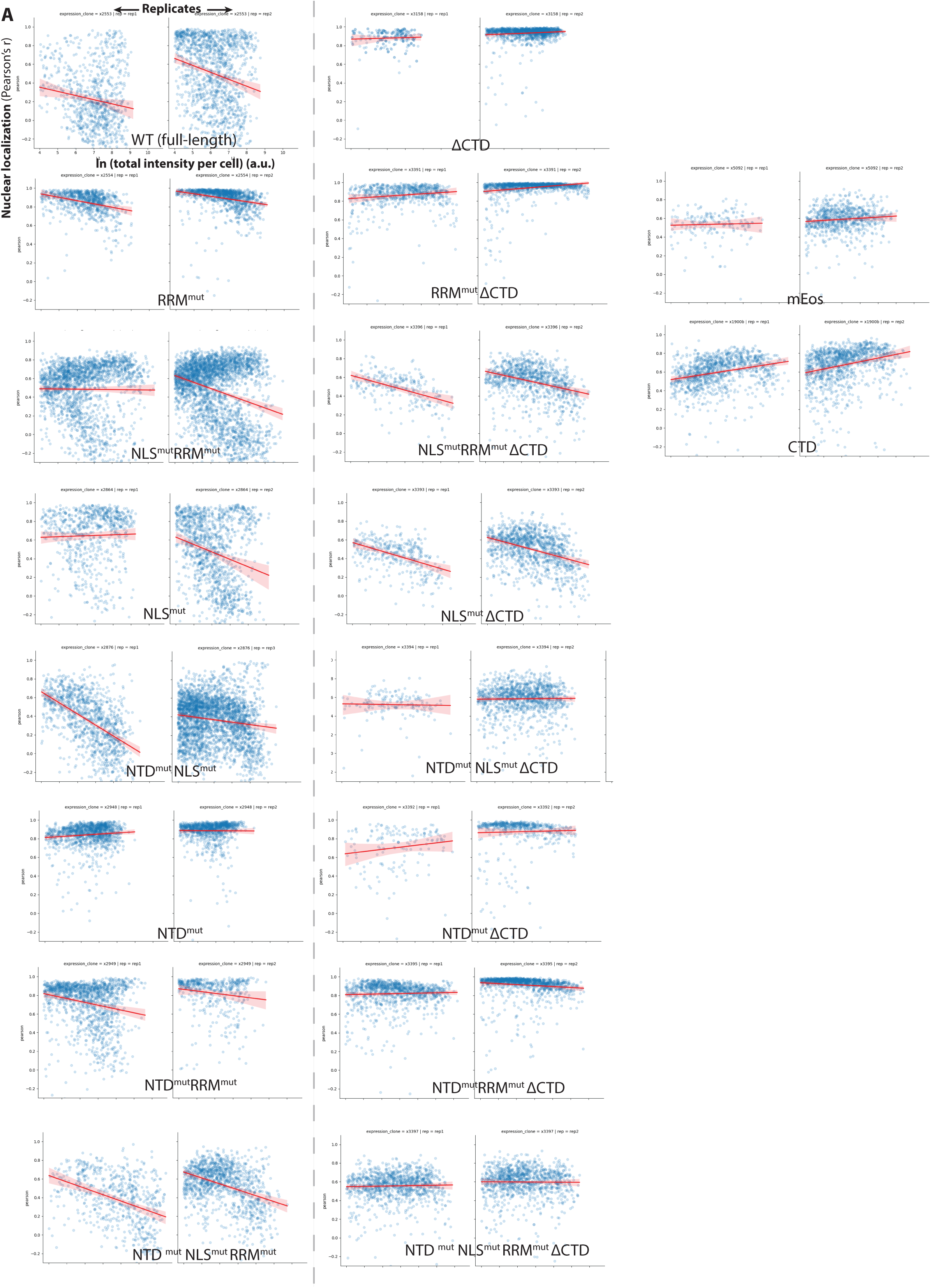
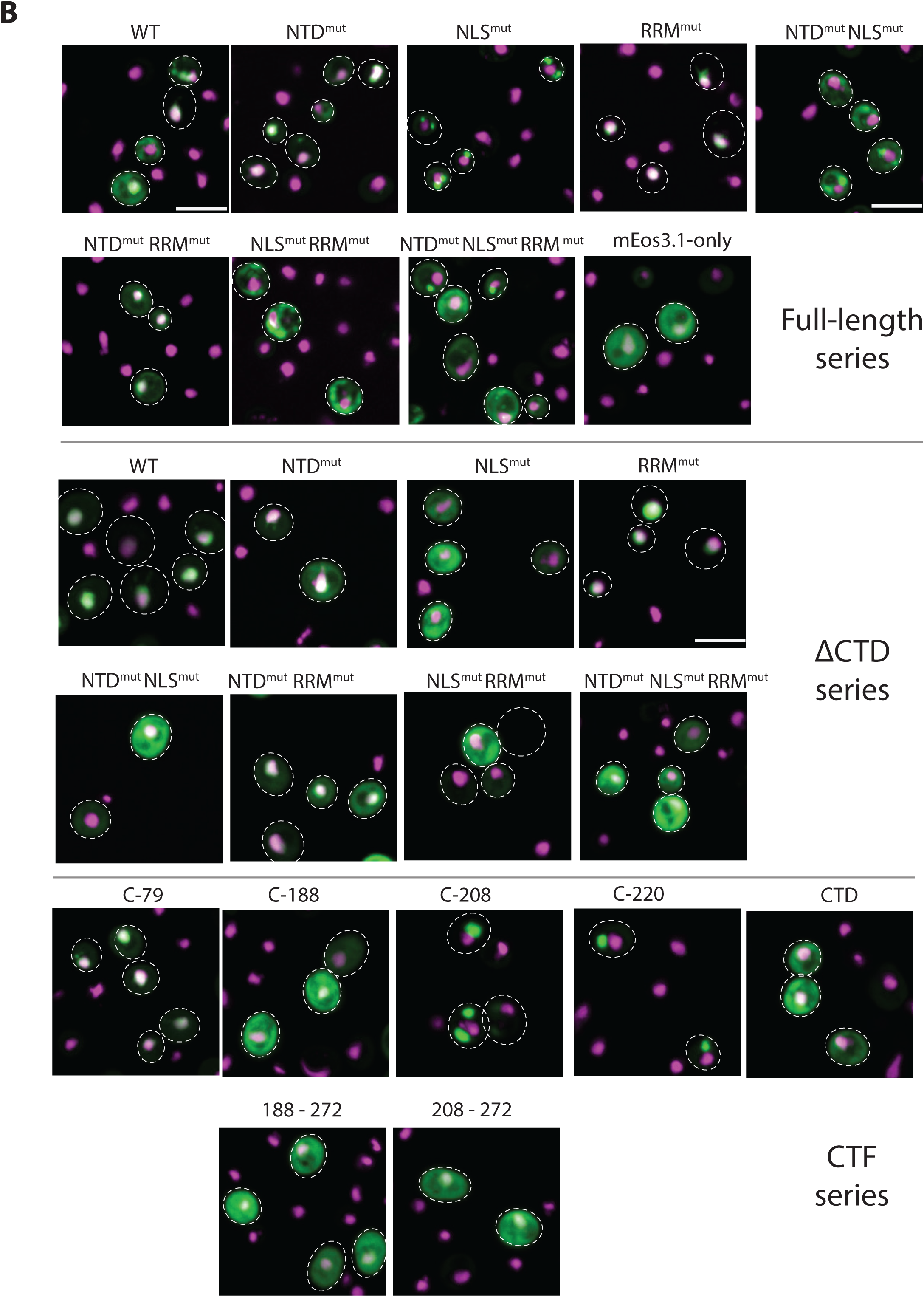
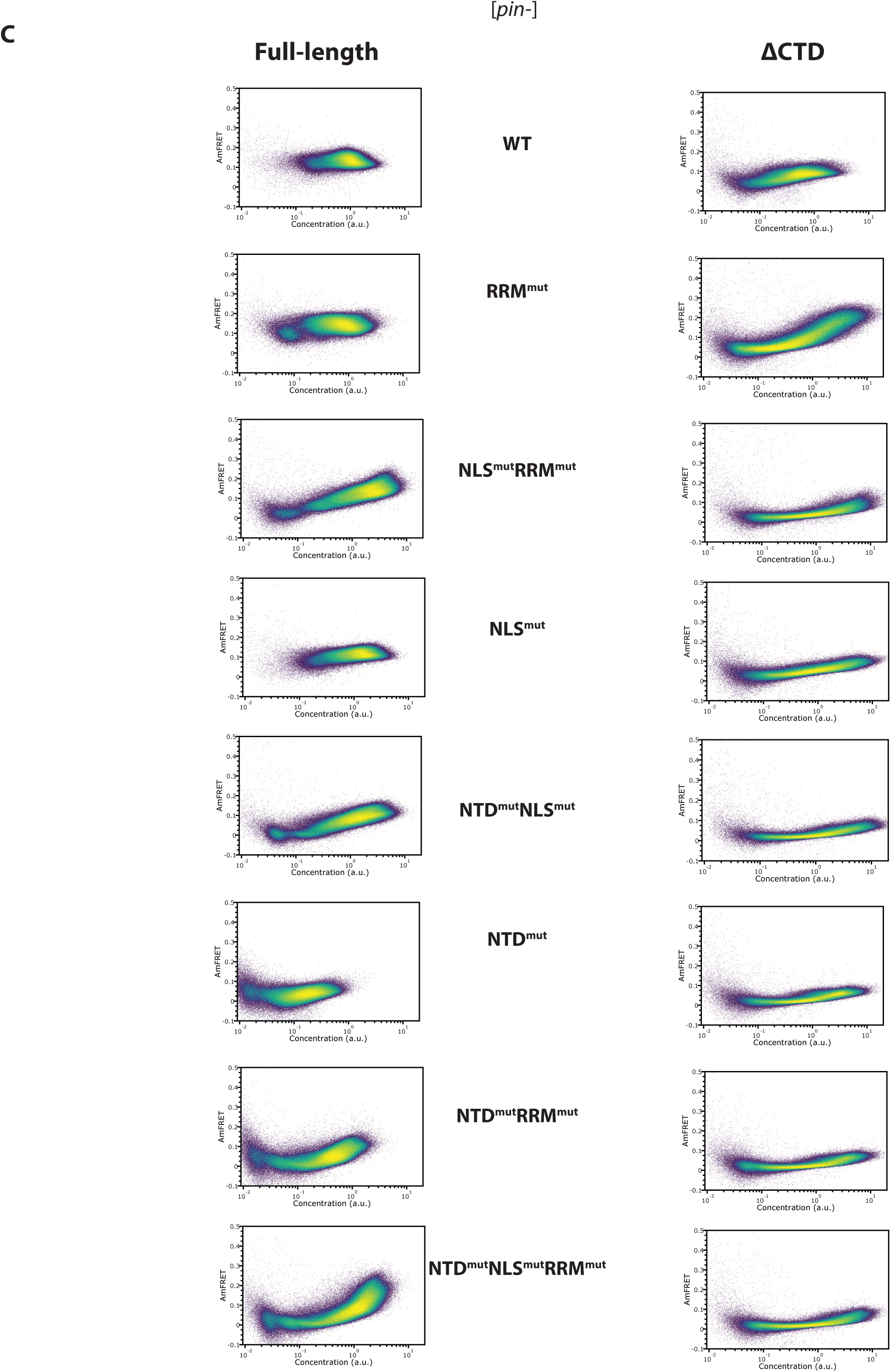
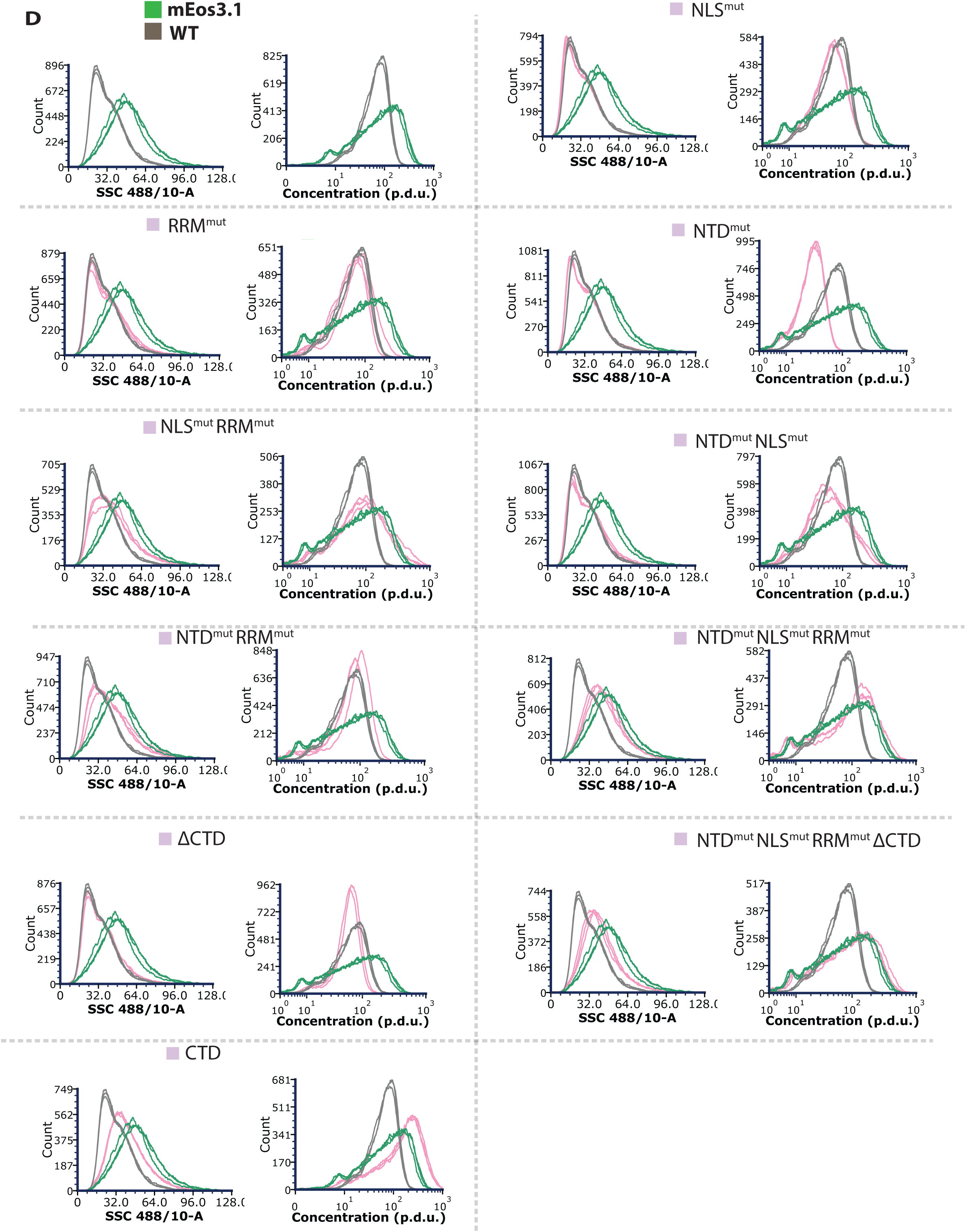
A) Scatter plot of single cell nuclear localization scores. (Pearson’s r) plotted against protein expression levels illuminate tight dependence of various mutants of TDP-43’s nuclear localization, cytosolic aggregation and protein expression. Population level expression appeared bimodal for all mutants, hence the cells with expression below ln(4) a.u. were gated out to exclude possible autofluorescence. B) **Representative montage** of max projected confocal imaging of a comprehensive list of full-length, ΔCTD and CTF constructs Note that cytoplasmic puncta in these [*pin^-^*] cells represent non-amyloid assemblies. Dotted circles/ellipses indicate the bright field cell outlines; each panel was picked to best represent the overall trend in a mutant’s behavior. Given the heterogeneous range of expression of various mutants, brightness was adjusted individually to depict the fluorescence according to expression levels. Only the cells with visually discernible fluorescence were marked by dotted circles. Protein marked by mEos (green), nucleus by HTB2-BDFP1.6:1.6 (magenta). Scale bars represent 10 μm. C) **The CTD is necessary for phase transitions.** Side-by-side comparison of DAmFRET of identical point mutants in the full-length versus the ΔCTD series. Related to main Fig. 1B. Plots representative of triplicates and multiple independent experiments. D) **TDP-43 expression induces a growth defect.** Flow cytometry read-out of cell-size parameter - SSC-A (side scatter) and side-scatter normalized protein expression (in this case non-photoconverted mEos to gain maximum resolution over reporting protein concentration) showing WT TDP-43 (grey traces) strongly reduces growth, compared to mEos-only (green traces). This defect is rescued in the NTD^mut^ RRM^mut^ background (blue), restoring a broader expression range. Systematic comparisons for all other full-length variants. These histogram overlays contain triplicates of a protein that are colored the same.

**Figure S2.**
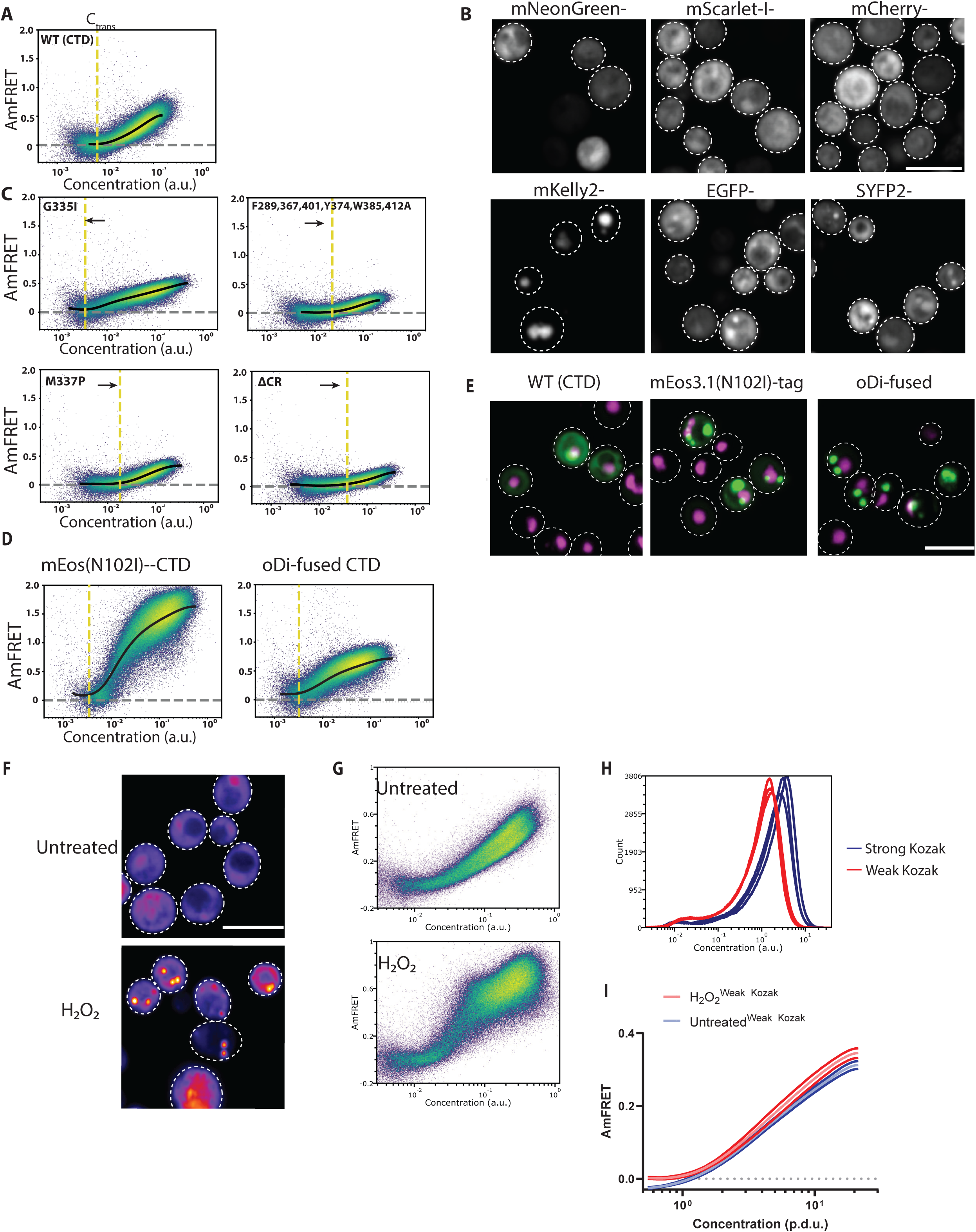
Characterization of CTD cluster stability, tag-dependency, and maturation. A) **CTD self-assembles cooperatively.** DAmFRET profile of mEos-CTD revealing monomer-like to non-stoichiometric, co-operative self-assembly. The yellow dashed line represents C_trans_ - an inflection point demarcating the onset of cooperative multimerization, derived from spline fitting (black trace), detailed in Methods. The grey dashed line represents median AmFRET of run-matched monomer-only mEos control via DAmFRET. B) A representative montage of hundreds of cells imaged with CTD tagged to commonly used fluorophores showing clear puncta with mKelly2, SYFP2 and EGFP, illustrating the dependence of CTD coalescence on fusion partner. Scale bar = 10 μm. C) DAmFRET and spline fit overlays of sequence mutants known to disrupt macromolecular phase separation of the CTD, showing they likewise govern subdiffraction clustering. Yellow dashes indicate the C_trans_ of the respective mutant and black arrows indicate the direction of change of C_trans_ with respect to CTD wild-type. D) DAmFRET plots with spline overlays of a dimerizing mutation to mEos (N102I) or a homodimer oDi, both of which induce a steep increase in AmFRET beyond C_trans_ while only shifting the baseline AmFRET level and apparent C_trans_ due to dimerization. E) Micrographs reveal condensation of the CTD as seen by large round puncta with depleted diffuse fluorescence, indicating that CTD clusters are normally arrested by internal saturation of interaction sites. Representative confocal max projected image of > 50 cells and duplicates at least; scale bar 10 μm. F) Microscopic validation of the exact treatment dose and time of H2O2 for the DAmFRET data collection - 8 mM for 2 hours, as opposed to the 3 mM dose for 6 hours on the time lapse imaging experiment (Fig. 2G). Scale bar = 10 μm. G) DAmFRET showing that H2O2 treatment as in F greatly increases AmFRET beyond C_trans_ without changing C_trans_ (shown in the shaded grey line covering both - the untreated and the treated plots.) H) Validation of the Kozak sequences working relative to each other - evidenced by a left-shifted range of protein concentration with the weak Kozak. I) Muted response of the CTD weak Kozak construct to H2O2, relative to that by the default, strong Kozak, suggesting a kinetic control of TDP-43 condensates.

**Figure S3.**
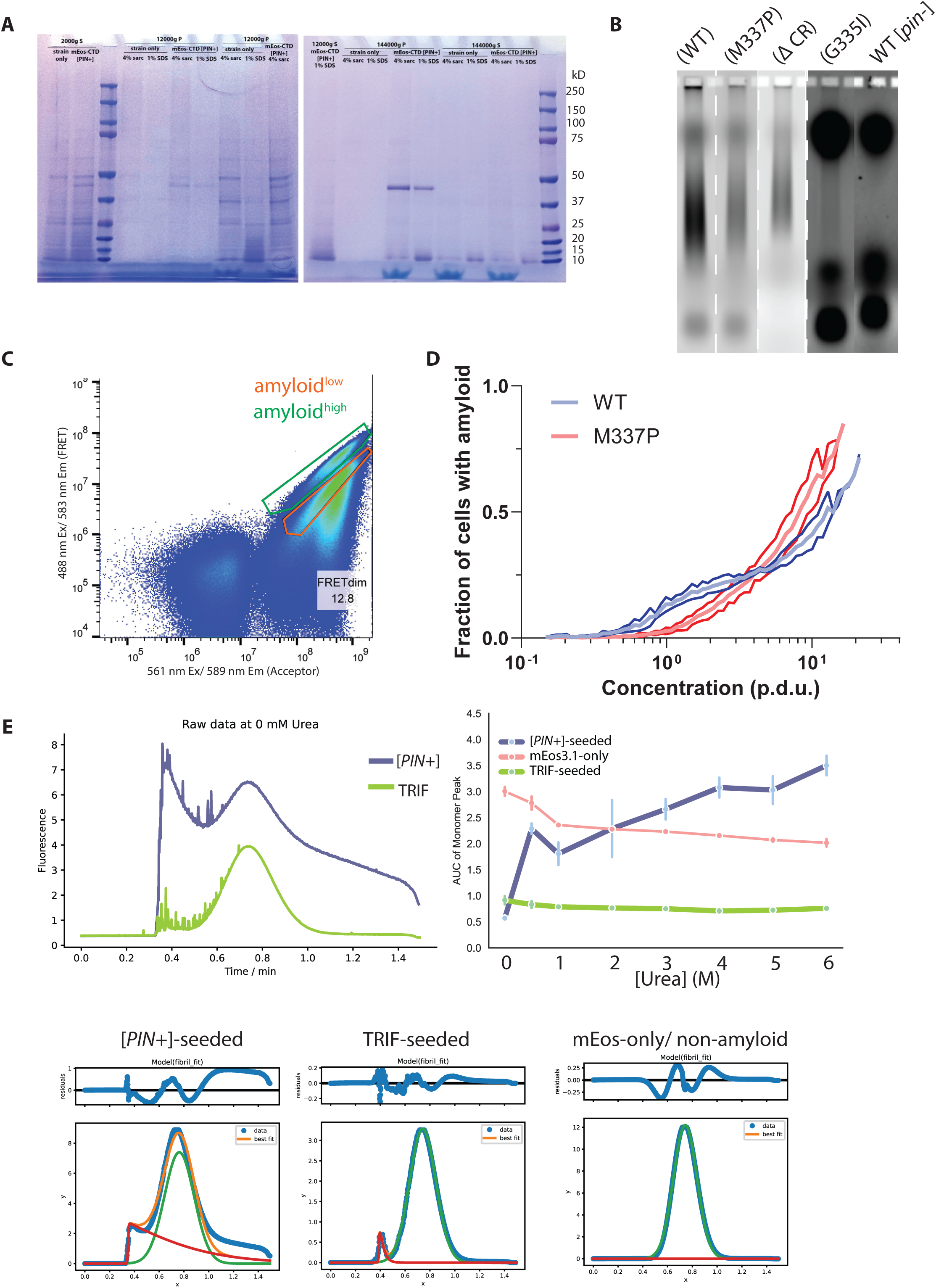
Characterization of amyloid kinetics, morphology, and stability. A) **Biochemical verification of amyloid and integrity of the fusion construct.** Coomassie-stained SDS-PAGE of the detergent-insoluble fraction from amyloid enriched lysates. A specific band corresponding to mEos-CTD appears only in [*PIN^+^*] cells, confirming the presence of insoluble aggregates. S = Supernatant fraction; P= Pellet fraction; Sarc = Sarkosyl; SDS = sodium dodecyl sulfate. B) **Extended biochemical validation for CTD amyloid using SDD-AGE.** WT CTD exhibited a characteristic, detergent resistant smear on agarose gel, specifically in [*PIN+*] but not in [*pin-*] cells. CTD mutations that reduce the fraction of amyloid-containing cells reduce the intensity of the smear. Brightness of [*pin-*] negative control adjusted to match G335I for accurate comparison. Dashed lines represent splicing of lanes from the same gel. Gel representative of multiple independent experiments. C) Gating of the two amyloid-containing populations of CTD-expressing cells used for FACS sorting, detailed in Methods. D) Overlaid WT and M337P CTD amyloid trace over the expression range, showing a steeper concentration-dependent increase in the fraction of amyloid post the C_trans_, for M337P. Black arrows indicate the direction of change of C_trans_ relative to WT CTD. E) Urea denaturation trace of amyloids measured by FIDA. (Left): Raw data of the Taylorgram showing generally more prodigious amyloid fraction in [*PIN^+^*]-seeded CTD than by TRIF. (Right): Area under the curve of monomer fraction of the lysate across all urea concentrations, showing modest and accountable amount of quenching of the fluorophore by urea. Fig. 3G is normalized according to this plot. (Bottom): Example from each of the samples showing the quality of fitting of the monomer Gaussian to the raw data.

**Figure S4.**
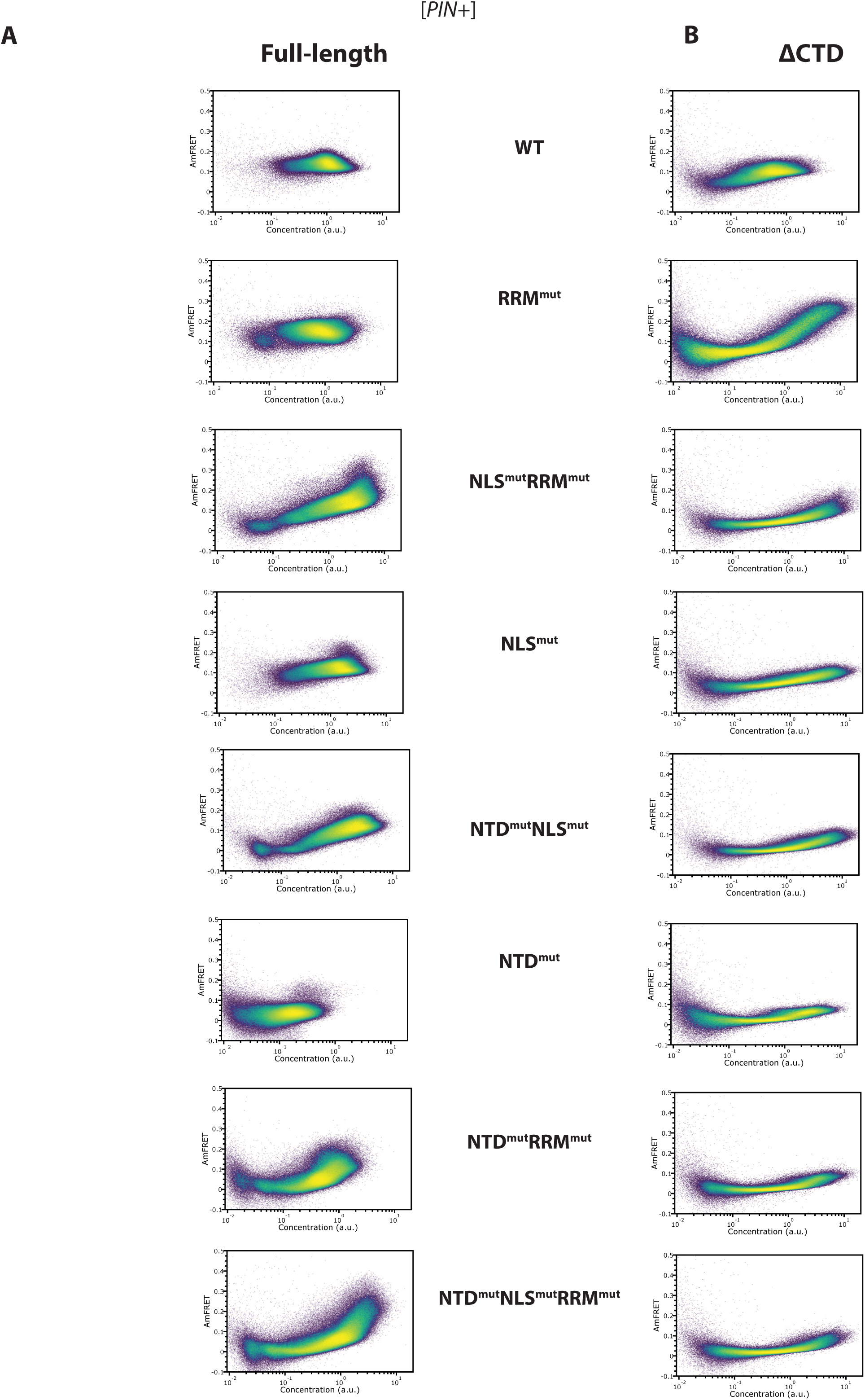
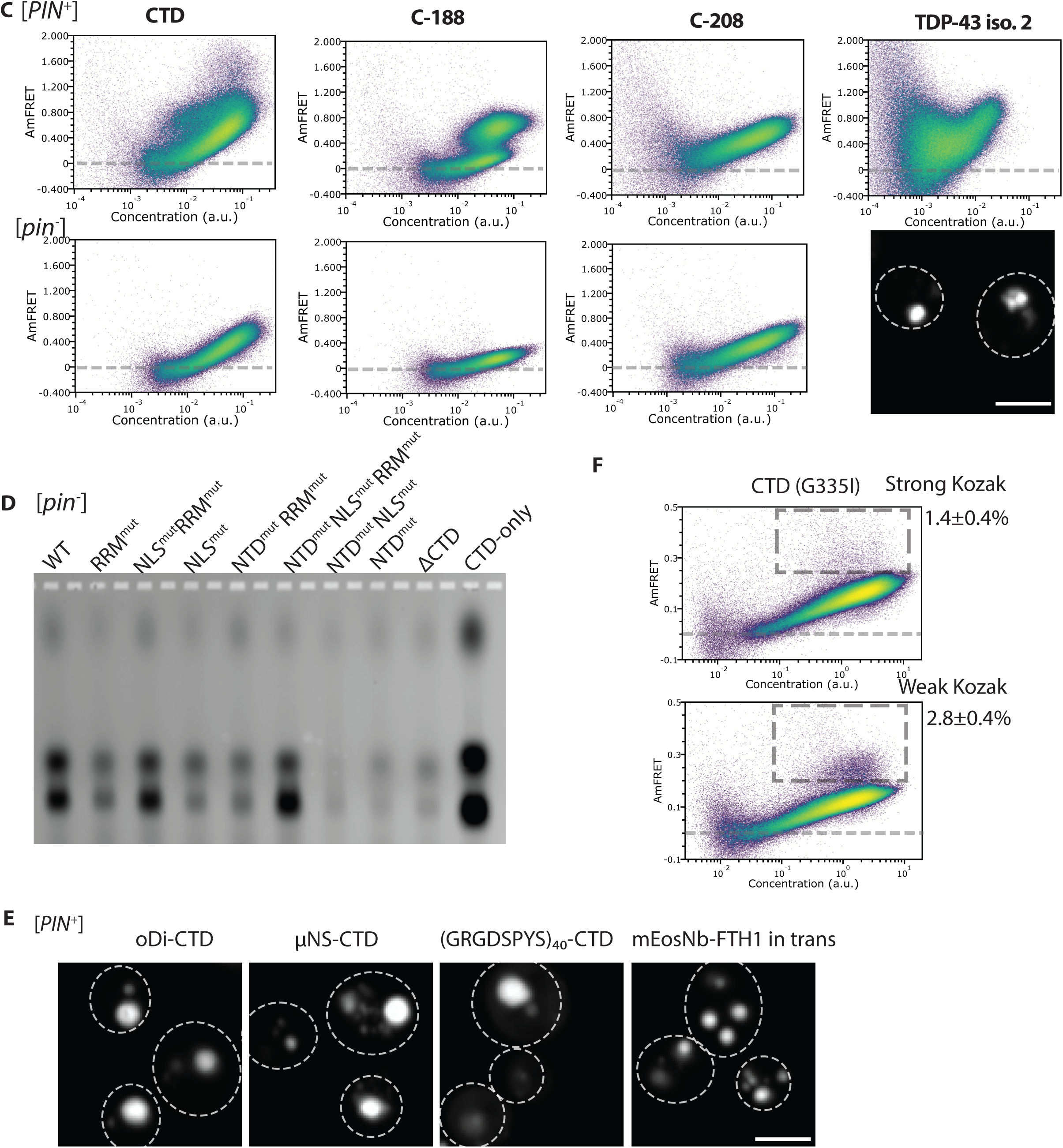
Extended characterization of amyloid suppression mechanisms. A) **Full DAmFRET profiles in [*PIN+*].** Extended data showing the representative DAmFRET profiles for all FL variants supporting the quantification in FIg. 4A. Notably, WT and RRM^mut^ in [*PIN^+^*] cells confirm their resistance to cross-seeding. In contrast, mutants involving the NLS and/or NTD show detectable amyloid formation. B) **Specificity of the amyloid formation.** DAmFRET profiles of ΔCTD variants in [*PIN^+^*] cells. Deletion of the CTD abolishes the high-AmFRET populations observed in NTD^mut^ and NLS^mut^ backgrounds, confirming that the amyloid signal is CTD-dependent. C) Side-by-side comparison of CTF DAmFRET profiles, showing that only CTD and C-188 have discontinuous populations characteristic of amyloid. Micrograph is a representative field of cells expressing TDP-43 isoform 2 exhibiting rampant aggregation. Scale bar = 5 μm. D) **SDD-AGE** in [*pin^-^*], corresponding to Figure 4B, showing that amyloid does not form in the absence of [*PIN*^+^] cross-seeding. E) **Microscopy of artificial condensates.** Representative image of cells expressing CTD fused to condensation-enhancing fusions. It drives the formation of large, spherical puncta, confirming they successfully drive phase separation despite inhibiting amyloid formation. Scale bar = 5 μm. F) **Flux-dependent amyloid prevalence.** Extended analysis of the Kozak effect, showing a de-repression of CTD (G335I) amyloid formation with slower translation flux.

